# An OSMR-CLIC1 cross talk drives key oncogenic pathways in glioblastoma

**DOI:** 10.1101/2025.03.05.641563

**Authors:** Amir Hossein Mansourabadi, Dianbo Qu, Francesca Cianci, Jamie Snider, Kamaldeep Randhawa, Laura Raco, Max Kotlyar, Mohammad Al Ayach, Guido Rey, Shridhar Sanghvi, Mark Abovsky, Harpreet Singh, Artee H. Luchman, Dylan Burger, Janusz Rak, Vahab D. Soleimani, Igor Jurisica, Igor Stagljar, Michele Mazzanti, Arezu Jahani-Asl

**Affiliations:** Department of Cellular and Molecular Medicine, Faculty of Medicine, University of Ottawa, 451 Smyth Road, K1H 8M5, Ottawa, Canada; University of Ottawa Brain and Mind Research Institute, 451 Smyth Road, K1H 8M5, Ottawa, Canada; Ottawa Institute of Systems Biology, 451 Smyth Road K1H 8M5, Ottawa, Canada; Laboratory of Cellular and Molecular Physiology, Department of Biosciences, University of Milano, Via Celoria 26, I-20133, Milano, Italy; Donnelly Centre, University of Toronto, Ontario, Canada; Division of Clinical and Translational Research, McGill University, 1001 Decarie Boulevard Montreal (Quebec) H4A 3J1; Osteoarthritis Research Program, Division of Orthopedic Surgery, Schroeder Arthritis Institute, and Data Science Discovery Centre for Chronic Diseases, Krembil Research Institute, University Health Network, 60 Leonard Avenue, 5KD-407, Toronto, Ontario, Canada M5T 0S8; Department of Physiology and Cell Biology, College of Medicine, The Ohio State University Wexner Medical Center, Columbus OH 43210; Department of Cell Biology and Anatomy, Arnie Charbonneau Cancer Institute | Hotchkiss Brain Institute, Cumming School of Medicine, University of Calgary, AB, T2N 4N1; Kidney Research Centre, Inflammation & Chronic Disease Program, Ottawa Hospital Research Institute, Ottawa, ON, K1H 8L6, Canada; Department of Pediatrics, Faculty of Medicine and Health Sciences, McGill University; Department of Biochemistry, Microbiology and Immunology, Faculty of Medicine, University of Ottawa, 451 Smyth Road K1H 8M5, Ottawa, Canada; Departments of Medical Biophysics and Computer Science, and Faculty of Dentistry, University of Toronto; Institute of Neuroimmunology, Slovak Academy of Sciences, Bratislava, Slovakia; Mediterranean Institute for Life Sciences, Meštrovićevo Šetalište 45, HR-21000 Split, Croatia; Department of Biochemistry, University of Toronto, Ontario, Canada; Department of Molecular Genetics, University of Toronto, Ontario, Canada; Regenerative Medicine Program and Cancer Therapeutics Program, Ottawa Hospital Research Institute, Ottawa, ON, K1H 8L6, Canada; Gerald Bronfman Department of Oncology, McGill University, 5100 de Maisonneuve Blvd. West, Montréal, QC, H4A 3T2, Canada

## Abstract

Oncostatin M receptor (OSMR) plays diverse and important roles in several human malignancies, including brain, breast, and pancreatic cancer.^1–4^ Glioblastoma (GB) is the most malignant genetically diverse brain tumour, with no cure. The most common genetic mutation in GB is a truncated active mutant of epidermal growth factor receptor (EGFR), the EGFRvIII. OSMR orchestrates a feedforward signaling mechanism with EGFRvIII and the signal transducer and activator of transcription 3 (STAT3), to drive GB progression.^4^ Beyond EGFRvIII, OSMR promotes brain tumour stem cells (BTSCs) via upregulation of mitochondrial oxidative phosphorylation and contributes to therapy resistance.^5^ The molecular mechanisms underlying the multifaceted roles of OSMR in different contexts are largely unclear. Here, we systematically mapped the OSMR interactome using Mammalian Membrane Two-Hybrid High-Throughput Screening (MaMTH-HTS). This unbiased approach led to the identification of OSMR-specific and OSMR/EGFRvIII-specific binding proteins, revealing context-dependent OSMR functions. Among a subset of common interactors, we uncovered chloride intracellular channel 1 (CLIC1) as a critical regulator of both OSMR-STAT3 signaling and the OSMR/EGFRvIII complex in GB. CLIC1 physically associates with both OSMR and EGFRvIII and plays a key role in EGFRvIII packaging into extracellular vesicles (EVs). Genetic deletion of CLIC1 disrupts the OSMR/EGFRvIII interaction, impairs STAT3 activation, reduces EGFRvIII EV content, and slows GB progression. Using whole-cell patch-clamp recordings and a monoclonal antibody that selectively targets transmembrane CLIC1 (tmCLIC1omab), we establish a distinct pharmacologically and biophysically tmCLIC-mediated current in GB indispensable for sustaining EGFRvIII/STAT3 signaling. Importantly, we show that OSMR is required for maintaining CLIC1-mediated ionic balance at the plasma membrane (PM). Our study uncovers a bidirectional crosstalk between OSMR and tmCLIC1 in GB, essential for fueling its malignant growth, and suggest that disrupting the OSMR/tmCLIC1 interaction provides a promising therapeutic avenue for GB treatment.

## Introduction

GB is the most aggressive incurable brain tumour with a diverse genetic profile. The oncogenic EGFRvIII mutant is present in more than 30% of GB patients and is initially assigned to the classical GB subtype. The mesenchymal GB subtype has also been described as one of the most malignant brain tumours. The aggressive nature of classical and mesenchymal subtypes is attributed in large part to the activity of STAT3, endowing the cells with the ability to enter an epithelial-to-mesenchymal (EMT)-like state, maintain stemness, grow, and metastasize.^6–9^ Recent studies have revealed that the cytokine receptor, OSMR, is a central player in propelling classical and mesenchymal GB subtypes. OSMR plays a crucial role in mediating oncogenic signal transduction initiated by the ligand OSM.^1,10^ The OSM-OSMR induces the activation of STAT3 as well as key oncogenic pathways including the phosphoinositide 3-kinase (PI3K)–AKT, c-Jun N-terminal kinases/mitogen-activated protein kinase (JNK/MAPK), and RAS/MAPK pathways.^11–13^ OSMR functions as a co-receptor for EGFRvIII in GB, inducing a positive feedback loop with STAT3.^4^ It also plays a crucial role in upregulating mitochondrial respiration, and conferring resistance to ionizing radiation (IR) therapy.^5^ Additionally, OSMR is a key regulator of brain tumour stem cell (BTSC) self-renewal^5^ and the immune microenvironment^14^, suggesting that its function is highly context-dependent.

Here, we employed MaMTH-HTS, a technology uniquely suited to identifying interacting partners of integral membrane proteins. Our screening revealed that OSMR physically and functionally interacts with tmCLIC1, positioning it as a central player in both OSM/OSMR cytokine signaling and the OSMR/EGFRvIII co-receptor complex. We demonstrate that the genetic deletion of tmCLIC1 and pharmacological inhibition of tmCLIC1 impairs BTSCs, and oncogenic EGFRvIII/STAT3 signaling. Furthermore, our data show that OSMR is essential for the function of the membrane configuration of CLIC1, highlighting a previously unrecognized crosstalk between a cytokine receptor and an ion channel in driving key oncogenic pathways in GB. Our finding has important implications for the design of novel therapeutic strategies to suppress GB.

## Results

### Mapping OSMR interactome via Mammalian Membrane Two-Hybrid High Throughput Screening (MaMTH-HTS)

To address the question of how OSMR networks with alternate molecules to confer its different functions, we performed MaMTH-HTS in HEK293T cells expressing OSMR bait, in the presence or absence of co-expressed EGFRvIII, alongside a prey library of ∼8000 open reading frames from the Human ORFeome V8.1 collection (**Supp Fig. 1a-c**).^15^ We identified 334 high-confidence candidate binding partners for OSMR (**Fig. 1**), following rigorous analyses and exclusion of MaMTH-HTS ‘frequent flyers’, which was established from repeated MaMTH-HTS results of 9 ‘individual bait’ protein control experiments, as previously described.^16^ Among the identified candidates, 204 hits, including 59 PM proteins, were detected when OSMR was used as bait in the absence of EGFRvIII, and 160 candidate binding partners, including 40 PM proteins, were identified when OSMR was used as a bait in the presence of EGFRvIII. Importantly, our data revealed 30 candidate binding partners of OSMR that were commonly shared by both groups, 7 of which were PM proteins. The PM proteins encompassed proteins that are integral to the membrane as well as proteins that are associated with the membrane. GO term functional analysis of protein clusters in each group revealed molecules engaged in metabolic processes and signalling as top categories underscored in each group (**Fig. 1**). Furthermore, these analyses revealed that the GO term for immune system processes was highly enriched in OSMR-unique interactome, whereas the regulation of biogenesis processes was among top GO terms for OSMR-EGFRvIII category. This novel OSMR interactome map highlights its role in different biological processes with unique variations depending on a cell’s genetic signature.

**Figure 1.**
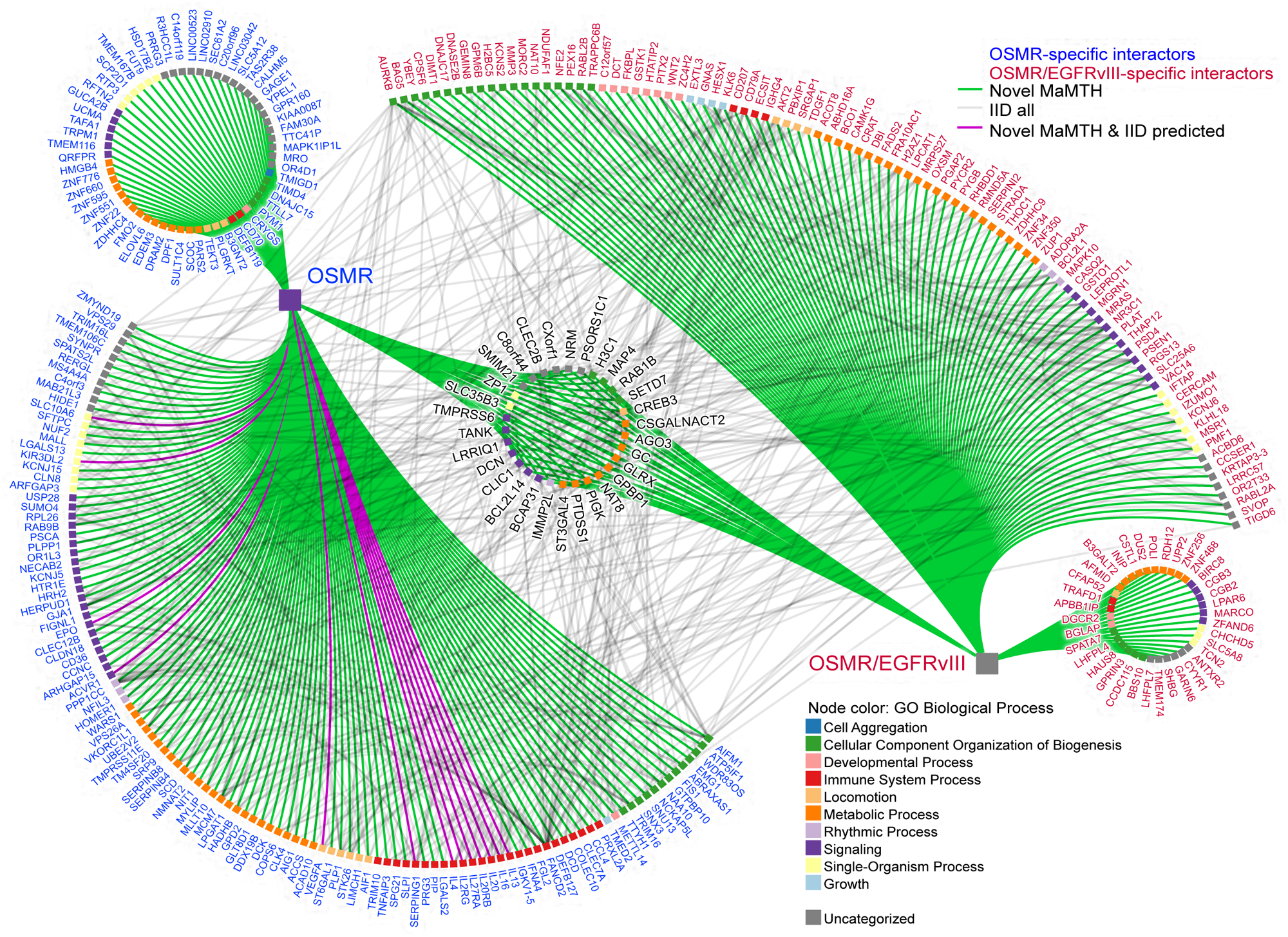
Mapping global binding partners of OSMR. Using MaMTH-HTS, we mapped the OSMR interactome in the absence and presence of EGFRvIII. The interactome is represented with green edges for the novel MaMTH candidates and gray edges highlighting known interactions identified using the Integrated Interaction Database (IID). Purple edges indicate interactions both identified in MaMTH and predicted by IID. The candidate binding partners of OSMR in the absence of EGFRvIII (OSMR-specific interactors) are represented in blue. The candidate binding partners of OSMR in the presence of EGFRvIII (OSMR/EGFRvIII-specific interactors) are represented in red. The candidate binding partners of OSMR that were commonly shared by both groups are shown in the center. Go term functional annotation of the potential candidates are shown with different coloured squares.

### Analysis of OSMR common binding partner and the involvement of CLIC1

To unravel the mechanism by which OSMR operates to promote tumour growth across different GB subtypes, we focused on the common candidate binding partners (**Supp. Table 1**), which encompassed different classes of proteins including metabolic processes, signalling, biogenesis, development, and immune system processes. To validate MaMTH data in the context of GB, we employed four different patient-derived BTSCs that naturally harbour EGFRvIII mutation (BTSC73 and BTSC147) or lack the mutation (BTSC12, BTSC30) (**Supp. Table 2**). We conducted a counter-screen in which we employed siRNA (**Supp. Table 3**) targeting each of the genes encoding these proteins, followed by assessing cell viability. Our results revealed that knockdown (KD) of 12 genes demonstrated a substantial reduction in cell viability exceeding 50% across different BTSCs (**Supp. Fig. 1d**). These included *CLEC2B*, *CLIC1*, *CREB3*, *CSGALNACT2*, *DCN*, *EIF2C3*, *GC*, *GPBP1*, *H3C1*, *IMMP2L*, *NAT8*, and *ST3GAL4*. Prior to follow up investigation on select candidates, we applied additional screening criteria pertaining to the known role of these proteins in BTSCs and GB as well as their known roles in regulation of different hallmarks of cancer. These criteria led us to focus on CLIC1, as the top candidate.

CLIC1 is highly expressed in various cancers including GB, with its expression significantly correlating with poor prognosis.^17^ CLIC1 exists as a soluble cytoplasmic protein, however, under specific cellular conditions that are favored in cancer cells^18^, it can translocate to PM where it functions as an ion channel.^19,20^ Interestingly, similar to OSMR^13^, CLIC1 is highly expressed in the mesenchymal GB subtype and enriched in cancer stem cells (CSCs)^17^, where it promotes proliferation.^21^ We conducted analysis of four publicly available patient datasets deposited in TCGA, Rembrandt, Gravendeel, Kamoun, and assessed the mRNA expression of CLIC1 and OSMR in GB tumour versus non-tumour samples. Both OSMR and CLIC1 were consistently upregulated in GB samples, across TCGA, Rembrandt, and Gravendeel datasets (**Fig. 2a,b**). Importantly, CLIC1 and OSMR expression analysis across different GB subtypes revealed that CLIC1 displays consistent and significant upregulation with subtype progression while OSMR was equally upregulated in both classical and mesenchymal subtypes relative to proneural subtype **(Fig. 2c,d**). Next, we analyzed how upregulation of OSMR and CLIC1 impacts GB patient survival. Analysis of 4 independent datasets deposited at TCGA, Rembrandt, Gravendeel and LeeY revealed that patients with high expression of CLIC1 or OSMR had the worst prognosis (**Supp. Fig. 2a-h**). Importantly, patients stratified as CLIC1/OSMR high expressors had significantly shorter lifespan compared to CLIC1/OSMR low expressors (**Fig. 2e-h**). Our data suggest that stratification of patients for CLIC1 and OSMR expression is an important predictor of survival.

**Figure 2.**
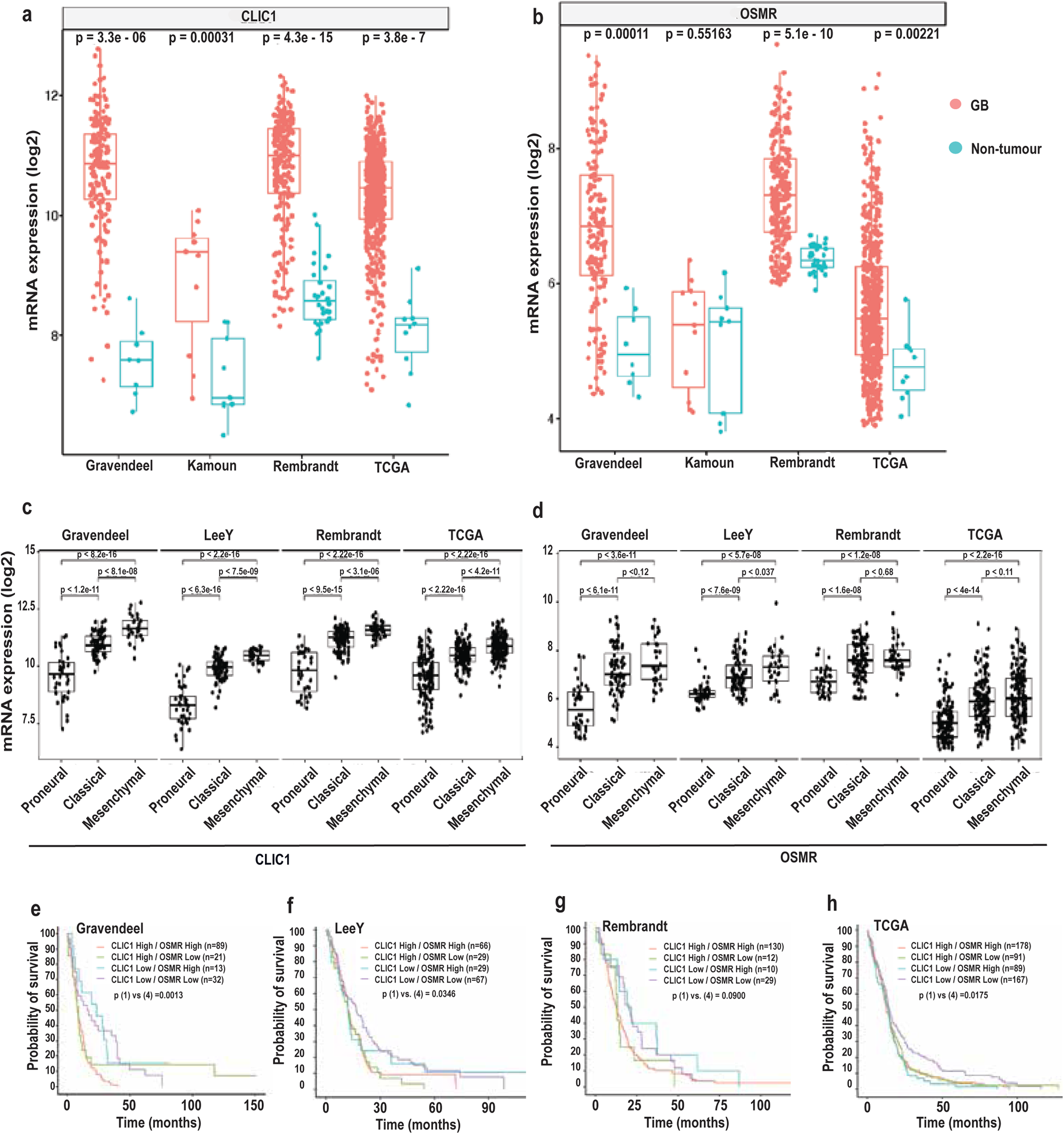
CLIC1 and OSMR are upregulated in glioblastoma, and their high expression correlates with poor patient survival. **a,b**, Expression of CLIC1 and OSMR in GB and non-tumour samples from four glioblastoma datasets (Gravendeel, Kamoun, Rembrandt, TCGA) was compared. Both genes were significantly elevated in GB samples, except for OSMR in the Kamoun dataset. **c,d**, Expression of CLIC1 and OSMR across different glioblastoma subtypes of Proneural, Classical, and Mesenchymal were ordered by disease severity in four datasets. CLIC1 displays consistent, significant upregulation with subtype progression, while OSMR is significantly upregulated in both Classical and Mesenchymal subtypes relative to proneural subtype. **e-h**, Clinical significance of variable CLIC1 expression in high- and low-OSMR expressing patients were analyzed using annotated public online datasets is shown.

### CLIC1 Regulation of BTSCs self-renewal and proliferation

To characterize the impact of CLIC1 knockdown on a panel of patient-derived BTSC with varying oncogenic mutations, we next electroporated a pool of four siRNAs targeting CLIC1 into different BTSCs and assessed cell viability and stem cell frequency using PrestoBlue and extreme limiting dilution assay (ELDA)^22^, respectively. RT-qPCR and immunoblotting analyses showed efficient siRNA-mediated knockdown of CLIC1 by greater than 85% and a significant decline in CLIC1 protein expression level relative to BTSCs electroporated with a non-targeting RNAi (siCTL) (**Fig. 3a-d, Supp. Fig. 3a-j**). Our data revealed a significant decrease in BTSC viability (**Supp. Fig. 3k,l**), BTSC sphere size (**Supp. Fig. 3m,n**), and stem cell frequency (**Fig. 3e-h**) upon KD of CLIC1. Next, we generated BTSCs in which we induced genetic deletion of CLIC1 using CRISPR in two different EGFRvIII-expressing BTSC73 and BTSC147 (**Fig. 3i-l**). Similar to our findings with the transient KD experiments, monoallelic (CRISPR-m) or biallelic (CRISPR-b) genetic deletion of CLIC1 induced a robust reduction in BTSC stem cell frequency (**Fig. 3m,n**), cell viability (**Supp. Fig. 3o,p**), and sphere size (**Supp. Fig. 3q,r**). We next conducted EdU assay. Consistent with our results, deletion of CLIC1 resulted in a significant reduction in actively dividing cells (**Fig. 3o,p, Supp. Fig. 4**). Taken together, our data confirm that CLIC1 is essential for maintaining cell viability, stem cell self-renewal and proliferation of BTSCs derived from glioblastoma patients.

**Figure 3.**
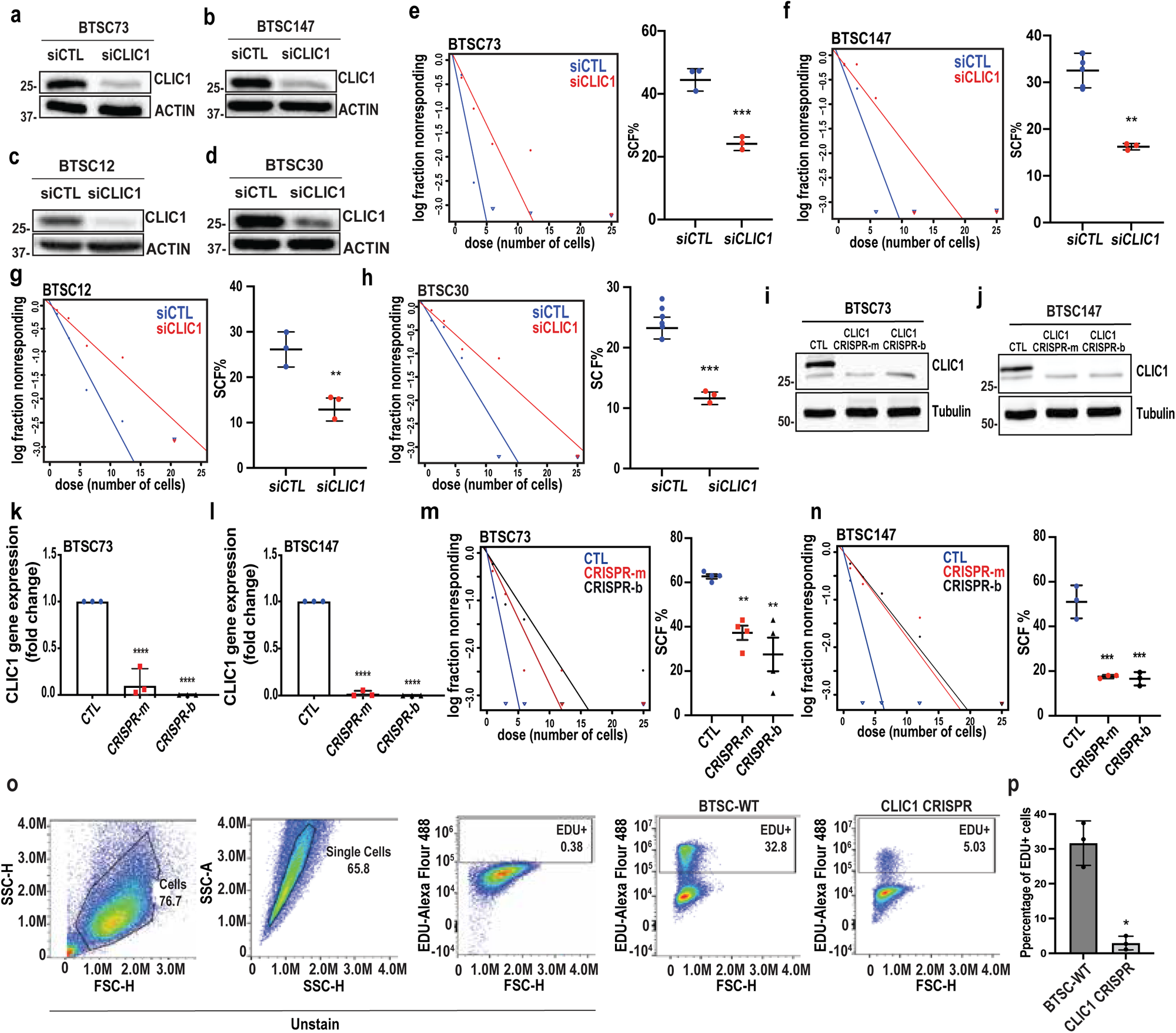
Cell viability evaluation following gene silencing of candidate binding protein targets. **a-d** Knockdown of CLIC1 was induced in BTSCs as described, and whole cell lysates were subjected to immunoblotting using a CLIC1 antibody. ACTIN was used as a loading control. **e-h**, CLIC1 knockdown and control cells were subjected to Extreme Limiting Dilution Assay, n=3 biological replicates. Statistical analysis was performed using a student-t test. Data are represented as the means ± SD. BTSC73 (**e**): \*\*\**P* = 0.0010; BTSC147 (**f**): \*\**P* = 0.0017; BTSC12 (**g**): \*\**P* = 0.0077; BTSC30 (**h**): \*\*\**P* = 0.0006 for each pairwise comparison. **i-j,** Validation of CRISPR-mediated CLIC1 KD and KO in patient-derived BTSC73 and BTSC147 cell lines. CLIC1 protein expression levels were evaluated by immunoblotting following CLIC1 KD (CLIC1 CRISPR-m) and CLIC1 KO (CLIC1 CRISPR-b), comparing them with the control group (CTL), with n=3 biological replicates. **k,l**, mRNA expression of CLIC1 was confirmed by RT-qPCR in CLIC1 CRISPR BTSCs 147 and 73. Gene expression was normalized to the housekeeping gene GUSB. Data for two different clones (CRISPR-m and CRISPR-b) are shown for each BTSCs and represented as the means ± SD, n = 3. Statistical analysis was performed using one-way ANOVA. BTSC73 (**k**): \*\*\*\**P* = 0.00003 and 0.00001, respectively, for each pairwise comparison from left to right. BTSC147 (**l**): \*\*\*\**P* = 0.000000006 and 0.000000005, respectively, from left to right; **m,n**, CLIC1 CRISPR and control BTSCs were subjected to ELDA and SCF was plotted. n=3 biological replicates/ Statistical analysis was preformed using one-way ANOVA. Data are represented as the means ± SD. BTSC73 CRISPR-m and CRISPR-b (**m**): \*\**P* = 0.0084, \*\**P* = 0.0011 for each pairwise comparison respectively. BTSC147 (**n**): \*\*\**P* = 0.0002; **o,** Flow cytometry analysis of DNA replication in EdU-labeled cells from BTSC73-CTL and BTSC73-CRISPR-m is shown. n=3 biological replicates. **p,** Statistical analysis was performed using a student-t test. Data are represented as the means ± SD, \**P* = 0.0107.

### CLIC1 interacts with OSMR and EGFRvIII endogenously in BTSCs

MaMTH data revealed that CLIC1 is a high confidence potential binding partner of OSMR. We first subjected different non-permeabilized BTSCs to immunostaining using an antibody to CLIC1 and observed localization of CLIC1 at the cell periphery (**Fig. 4a-c**). To assess whether CLIC1 and OSMR colocalize in BTSCs, we conducted co-immunostaining using antibodies to OSMR and CLIC1. Our data revealed that CLIC1 colocalizes with OSMR in all the BTSCs analyzed in our studies (**Fig. 4d-g**), raising the question of whether CLIC1 interacts with OSMR endogenously. We conducted Proximity Ligation Assay (PLA) using OSMR and CLIC1 antibodies in different BTSCs and used BTSCs subjected to PLA with primary antibodies omitted as control. Our data showed *in situ* interaction of CLIC1 with OSMR across all BTSC lines that were examined (**Fig. 4h-l**), with EGFRvIII-expressing BTSC 73 and 147 that harbour elevated OSMR expression, exhibiting stronger signal (**Fig. 4h,i, l**) compared to BTSCs 30 and 12 lacking the EGFRvIII (**Fig. 4j,k, l**). To further confirm the specificity of the interaction signal, we subjected CLIC1 CRISPR-BTSCs and their corresponding controls to PLA analysis. For each BTSCs two CRISPR lines, CRISPR-m and CRISPR-b, were included. Our results showed that the PLA interaction signal of OSMR-CLIC1 was robustly diminished in CLIC1 CRISPR BTSCs compared to control BTSCs (**Fig. 4m-p**).

**Figure 4.**
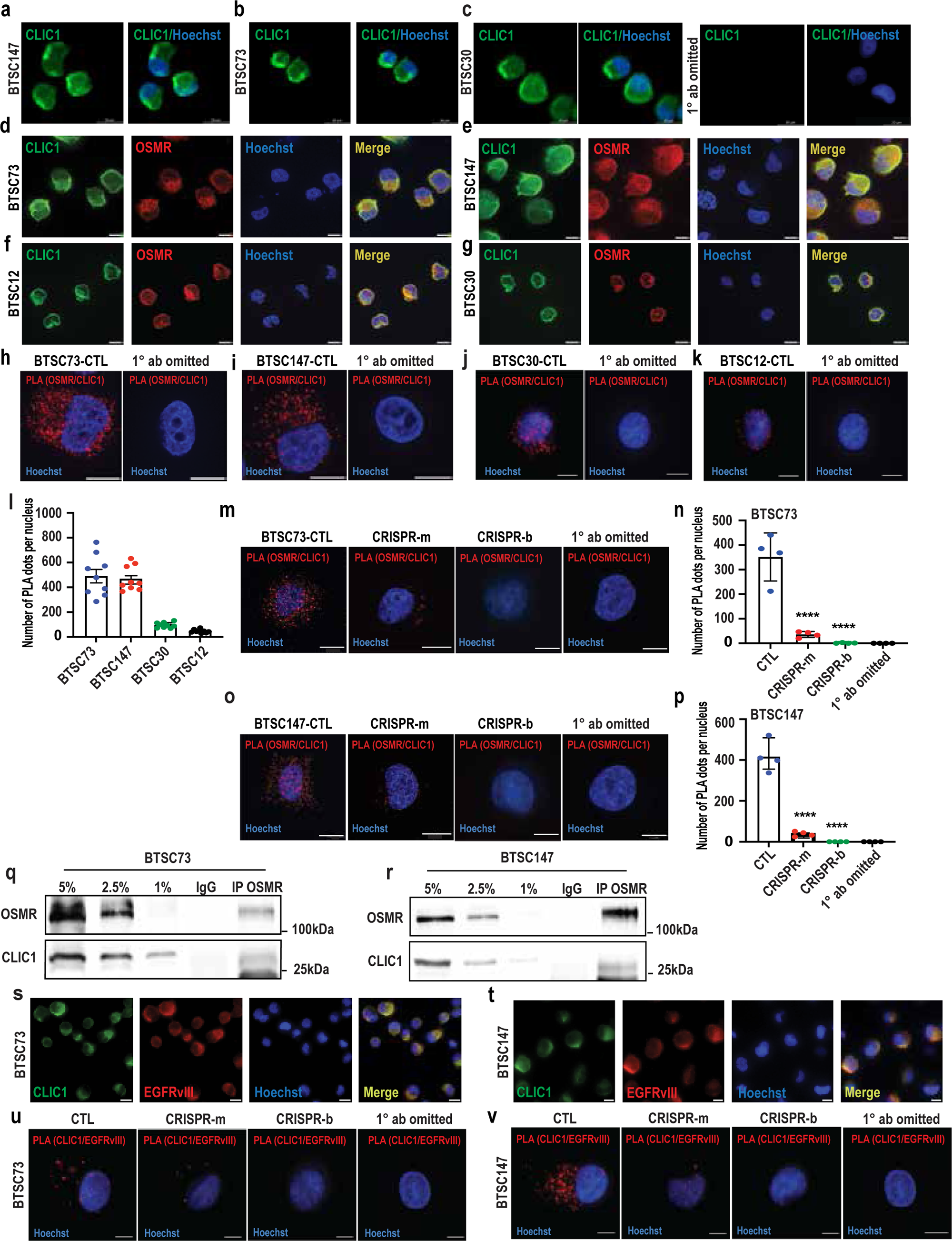
CLIC1 and OSMR interact endogenously in BTSCs. **a-c**, Non-permeabilized BTSCs were subject to immunostaining and CLIC1 is detected at the plasma membrane. Images in Merged shows representative CLIC1 expression (green) overlapping with Hoechst (blue). Each experiment was conducted with a primary antibody omitted control group, n=3 biological replicates. **d-g**, BTSCs were subjected to co-immunostaining using antibodies to CLIC1 (green) and OSMR (red). Nuclei were stained with Hoechst. Merged images depict co-localization of OSMR and CLIC1 (yellow). n=3 biological replicates. **h-l**, BTSCs were subjected to proximity ligation assay (PLA) using antibodies to CLIC1 and OSMR. Each red puncta represents PLA signal. Primary antibodies were omitted for control group. Nuclei were stained with Hoechst. **m-p**, The interaction of CLIC1 and OSMR was assessed by PLA in CLIC1 CRISPR and control BTSCs as described in h-l. Nuclei were stained with Hoechst. The number of PLA puncta (red) per nucleus was quantified in CRISPR and control BTSC73 and BTSC147 (**n,p**). Data are presented as the means ± SEM. Each dot represents the number of PLA dots per nucleus. A minimum of n=3 biological replicate images was assessed for statistical analysis. Statistical analysis was performed using one-way ANOVA. BTSC73 (**n**): ****P =0.000002 and 0.0000009 respectively, for each pairwise comparison from left to right. Scale bar, 10 μm; BTSC147 (**p**): ****P = 0.00000003 and 0.00000001 respectively, from left to right. **q,r**, BTSCs 73 or 147 were subjected to immunoprecipitation (IP) using an OSMR antibody or IgG control and pull downs were analyzed by immunoblotting using CLIC1 antibody. **s,t**, BTSCs 73 and 147 were subjected to co-immunostaining using antibodies to CLIC1 and EGFRvIII. Representative images represent n=3 independent biological replicates. **u,v**, CLIC1 CRISPR and control BTSCs were subjected to PLA analysis using antibodies to CLIC1 and EGFRvIII.

Next, we subjected the BTSCs to co-immunoprecipitation (co-IP) experiments using an antibody against endogenous OSMR followed by immunoblotting with a CLIC1 antibody. Our data showed that OSMR interacts with CLIC1 endogenously in BTSCs (**Fig. 4q,r**).

Given that OSMR forms a co-receptor with EGFRvIII, we next asked whether CLIC1 is a component of the same complex with EGFRvIII. We conducted co-immunostaining analysis in different BTSCs and found colocalization of EGFRvIII with CLIC1 (**Fig. 4s,t**). Thus, we performed PLA experiments using antibodies to EGFRvIII and CLIC1 in control and CLIC1 CRISPR BTSCs. Our results showed *in situ* interaction of CLIC1 with EGFRvIII (**Fig. 4u,v**).

### The extracellular domain of CLIC1 interacts with OSMR/EGFRvIII

OSMR is found in complex with EGFRvIII to amplify receptor tyrosine kinase (RTK) signalling.^4^ Given our results that CLIC1 interacts with both OSMR and EGFRvIII, we asked whether CLIC1 is required for the OSMR/EGFRvIII interaction. We subjected CLIC1 CRISPR and corresponding control BTSCs to PLA analysis using antibodies to OSMR and EGFRvIII. Strikingly, genetic deletion of CLIC1 significantly impaired the interaction of OSMR with EGFRvIII in each of BTSC147 and BTSC73 that harbour EGFRvIII mutation (**Fig. 5a-e**), revealing that CLIC1 is required for OSMR/EGFRvIII complex. In parallel, we conducted similar experiments in which we assessed CLIC1-OSMR interaction in EGFRvIII knockdown BTSCs and EGFRvIII-CLIC1 interaction in OSMR knockdown BTSCs. Our results suggest that CLIC1 and OSMR interact independently of EGFRvIII, although the presence of EGFRvIII generates stronger interaction signal (**Supp. Fig. 5a**), most probably due to the fact that OSMR expression is upregulated downstream of EGFRvIII-STAT3 signalling.^4^

**Figure 5.**
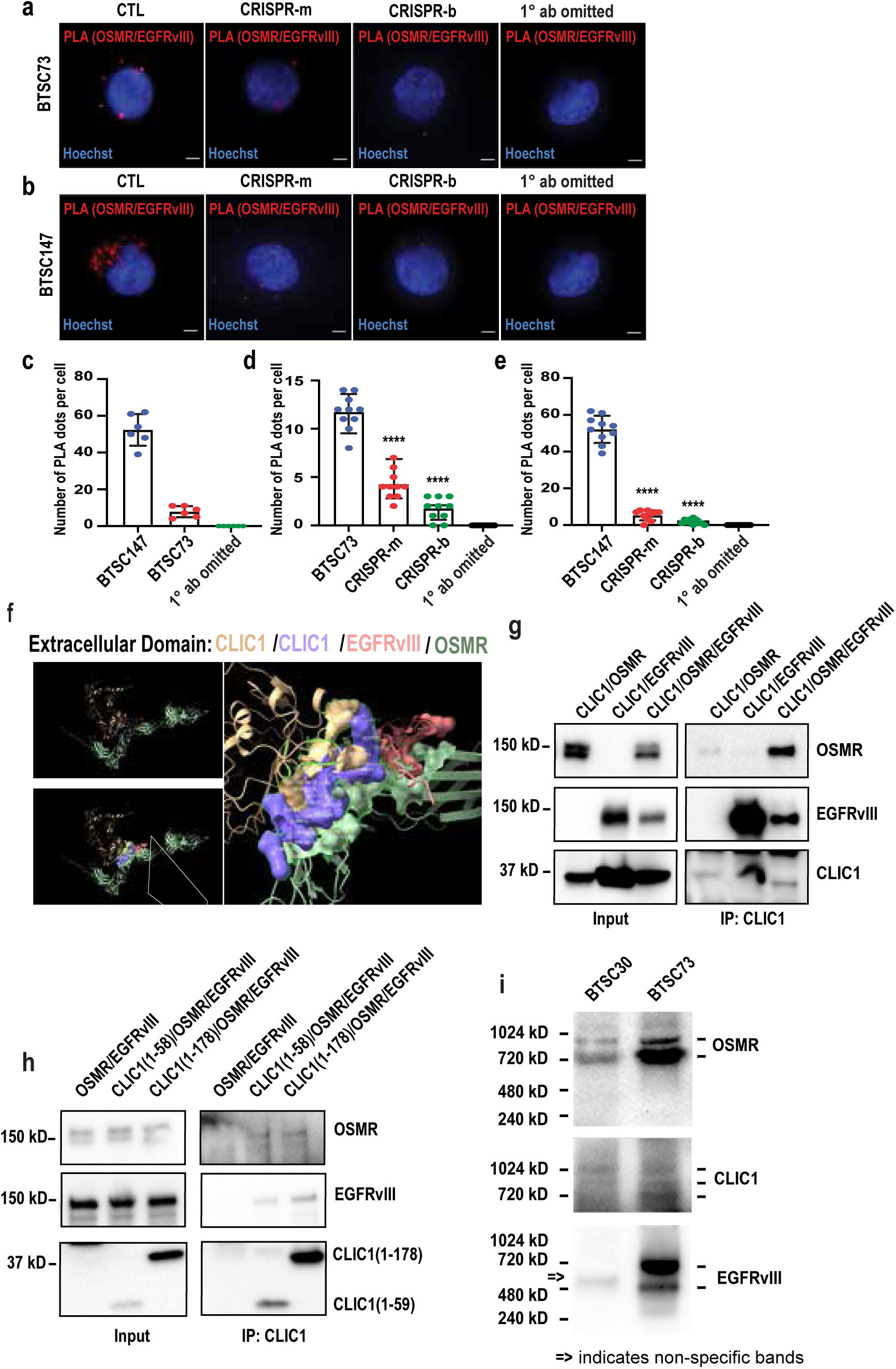
CLIC1 mediates the formation of the OSMR/EGFRvIII complex through its extracellular domain. **a-e** BTSCs 73, 147, and their related CLIC1 CRISPR cells were subjected to PLA analysis using antibodies to OSMR and EGFRvIII (**a,b**) and quantification of # of PLA per nuclei are presented (**c-e**). Nuclei were stained with Hoechst. Scale bar, 10 μm. Statistical analysis was performed using one-way ANOVA. BTSC73 (**d**): \*\*\*\**P* = 0.000000001 and 1 × 10^−13^ respectively, from left to right; BTSC147 (**e**): \*\*\*\**P* = 1 × 10^−15^ for each pairwise comparison. **f**, The interaction of the CLIC1 dimer (purple and brown), OSMR (green), and EGFRvIII (orange) was predicted using AlphaFold-Multimer for their extracellular domains. The predicted 3D structures of the extracellular domains, including surface representations of the interaction regions and close-up views of the binding interfaces between CLIC1 and EGFRvIII, and between EGFRvIII and OSMR, are shown. **g,h,** Experimental validation of the predicted domain using co-IP was performed. HEK293T cells were transfected with plasmids encoding CLIC1, OSMR, and/or EGFRvIII. Protein extracts were subjected to CLIC1 IP, followed by WB detection of CLIC1, OSMR, and EGFRvIII (**g**). HEK293T cells were transfected with plasmids encoding truncated CLIC1 constructs, CLIC1(1–58) and CLIC1(1–178), together with OSMR and/or EGFRvIII. Protein extracts were subjected to CLIC1 IP, and CLIC1, OSMR, and EGFRvIII were detected by WB (**h**). **i**, Native gel electrophoresis for CLIC1, OSMR, and EGFRvIII is shown.

Given that CLIC1 functions as a membrane protein existing predominantly as a homodimer, we next modeled its interaction using the dimer of CLIC1(residues 1-24) extracellular domain with the extracellular domains of OSMR (residues 1–741) and EGFRvIII (residues 1–378) using AlphaFold-Multimer. The predicted structural model of this tetramer revealed that the CLIC1 dimer (purple and brown) interacts with OSMR (green) and EGFRvIII (orange) (**Fig. 5f**). Given that the extracellular region of CLIC1 comprises only 24 amino acids (aa), we included its transmembrane domain (residues 25–58 aa) for binding assays. Consistent with the prediction, our experiments using CLIC1(1–58 aa) and CLIC1(1–178 aa) as baits demonstrated that CLIC1(1–58 aa) interacts with OSMR or with the OSMR/EGFRvIII complex, indicating that the extracellular regions of these three proteins are sufficient to support complex formation (**Fig. g,h**). To further validate the interaction under native conditions, we performed native gel electrophoresis, which preserves intact protein complexes. The results revealed that CLIC1, OSMR, and EGFRvIII coexist within a ∼720 kDa protein complex in EGFRvIII-expressing BTSCs (**Fig. 5i**). Additionally, we confirmed that the protein expression of each of the molecules in the complex is not altered by the presence of its binding partners as revealed via immunoblotting analysis in BTSCs (**Supp Fig. 5b-d**), and IF staining on tumour tissues (**Supp Fig. 5e**). To investigate the regulatory relationship between EGFRvIII and STAT3 in controlling CLIC1 expression, we used EGFRvIII-expressing STAT3^loxp/loxp^, STAT3^−/-^, murine stem cell virus (MSCV)-CTL (non– EGFRvIII-expressing) STAT3^loxp/loxp^ and STAT3^−/-^ astrocytes. No differences in CLIC1 expression were observed among the groups, indicating that STAT3 does not regulate CLIC1 expression (**Supp Fig. 5f**). Next, we asked if the complex formation is impacted by lipid rafts bringing these molecules together. We treated BTSCs with Methyl-β-cyclodextrin (MβCD), to disrupt lipid rafts, followed by PLA analysis. We observed a reduction in the interaction signal between CLIC1 and OSMR suggesting that their interaction in part occurs within these lipid rafts on the plasma membrane (**Supp Fig. 6**).

**Figure 6.**
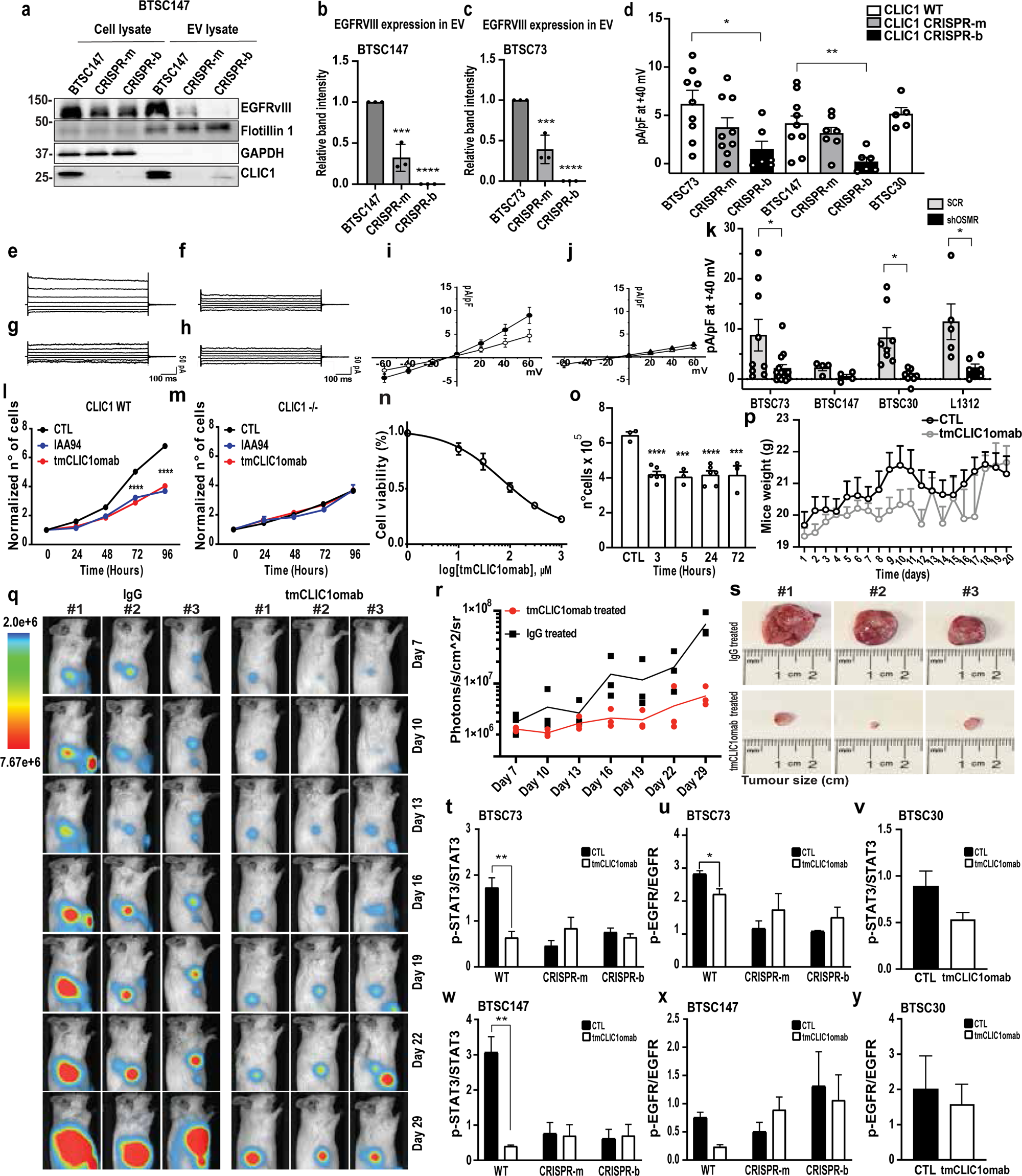
Blocking of tmCLIC1 using a monoclonal antibody reduces tumourigenesis, decreases tumour size, and suppresses STAT3 phosphorylation. **a-c,** CLIC1 CRISPR and control BTSCs were subjected to extracellular vesicle (EV) isolation. EV-digested lysates and whole cell lysates were subjected to immunoblotting analysis using antibodies indicated on the blot (**a**). Quantification of EGFRvIII in EV were conducted (**b,c**). Statistical analysis was performed using one-way ANOVA. n=3 biological replicates. BTSC147 (**b**): \*\*\**P* = 0.00025, \*\*\*\**P* = 0.000027; BTSC73 (**c**): \*\*\**P* = 0.00098, \*\*\*\**P* = 0.000059 for each pairwise comparison. **d**, Electrophysiological evaluation of tmCLIC1 was conducted at +40 mV on CLIC1 CRIPSR and control BTSCs. WT (white bar), CLIC1 +/- (monoallelic deletion, grey bar) and CLIC1 -/- (biallelic deletion, black bar). \**P* = 0.024, \*\**P* = 0.0091. **e-h**, Representative whole cell current for cells were conducted at several voltages ranging from -60 to +60 mV with 20 mV increments in control (**e,g**) and in OSMR knockdown BTSCs (**f,h**). Currents were recorded in resting conditions (**e,f**) and after IAA94 administration (**g,h**). **i**, I/V plot for control (SCR, black circles) and after perfusion of IAA94 (empty circles), and **j**, I/V plot for shOSMR cells in CTL condition (black triangles) and after perfusion of IAA94 (empty triangles) are shown. Average current-voltage recordings at different test potentials are shown. **k**, tmCLIC1 current for four different GB cell lines were analyzed at +40 mV in shOSMR and SCR control groups. The average current for the SCR (grey) and shOSMR (black) cells are shown. BTSC73 \**P* = 0.0179; BTSC30 \**P* = 0.0173, L1312 \**P* = 0.0205. **l,m**, Growth curves of human primary GB cell lines, WT and CLIC1 knockout, are shown. (**l**) GB human primary cell line WT growth curves over 96 hours in the absence (black circles) or presence (red circles) of tmCLIC1omab and IAA94 (blue circles); CTL, 24-96 hours, n=4; tmCLIC1omab and IAA94, 24-96 hours, n=7. 72-96 hours, ****p<0.0001. Means ± SEM, two-way ANOVA, Tukey’s multiple comparison test. (**m**) GB human primary cell line knockout for CLIC1 protein growth curves over 96 hours in the absence (black circles) or presence (red circles) of tmCLIC1omab and IAA94 (blue circles). CTL, 24-96 hours, n=5; tmCLIC1omab and IAA94, 24-96 hours, n=6. Means ± SEM. **n**, Dose-response curve of human primary GB cell line treated with increasing concentrations of tmCLIC1omab is shown. Cell viability was measured after 72 hours of treatment. The response was fitted using a non-linear regression model (four-parameter logistic model), and IC₅₀ (half-maximal inhibitory concentration) was calculated. Error bars represent the standard deviation (SD) of 4 biological replicates. IC50=0.184 µM. **o**, tmCLIC1omab effect on proliferation after several time of incubation is shown. Cell number of human primary GB cells incubated for 3, 5, 24 or 72 hours with tmCLIC1omab. The antibody was washout after selected time of incubation and then the number of cells was evaluated after 72 hours. One-way ANOVA; CTL (n=3) vs 3h (n=6) ****p<0,0001; CTL vs 5h (n=3) *** p=0,0002; CTL vs 24 h (n=6) **** p<0,0001; CTL vs 72h (n=3) *** p=0,0003. Tukey’s multiple comparison test. **p**, Weight of the animals treated or not treated with tmCLIC1omAb is shown. Body weight was measured prior to each injection and is presented as mean ± SEM (n = 8 mice per group). No significant weight loss was observed over the treatment period, indicating that tmCLIC1omab was well tolerated. Statistical significance was assessed using two-way ANOVA, Sidak’s multiple comparison test. **q-s,** BTSC73 cells were xenografted subcutaneously into 8-week-old SCID male mice, and treatment with 200 µg tmCLIC1omAb per animal, three times a week, was started on day 7 post-tumour inoculation. IVIS imaging was conducted to evaluate tumour volume (**q**), with the signal quantified (**r**). The tumours were then resected from the animals, and tumour size was measured using a standard method (**s**). **t-y**, CLIC1 CRISPR and control BTSCs were treated with tmCLIC1omab for 72 hours and were analyzed by immunoblotting using antibodies to p-STAT3, STAT3, p-EGFR and EGFR. Densitometric analysis of p-STAT3 to total STAT3 **(t,v,w**) and p-EGFR to total EGFR (**u,x,y**) in tmCLIC1mab treated antibodies versus control are shown. Statistical analysis was performed using unpaired students’ t test. n = 3 or > 3 biological replicates. BTSC73 (**t**): \*\**P* = 0.0022; BTSC73 (**u**): \**P* = 0.0335; BTSC147 (**w**): \*\**P* = 0.0038.

### OSMR-CLIC1 cross-talk maintain EGFRvIII/STAT3 signalling, and blockade of transmembrane CLIC1 (tmCLIC1) impairs the phosphorylation of EGFRvIII and STAT3

Our data raised the question of whether CLIC1 functionally regulates the oncogenic OSMR-STAT3 and EGFRvIII signalling. CLIC1 is found as a soluble cytoplasmic protein as well as a transmembrane protein that forms an ion channel.^21^ We conducted cell fractionation experiments in different BTSCs and found that CLIC1 is localized to both the membrane and cytoplasm (**Supp. Fig. 7**), although, the majority of CLIC1 was found in the cytoplasm. CLIC1 is shown to be packaged into extracellular vesicles (EVs) in order to target neighbouring cells^23^, promoting tumour growth.^23,24^ Interestingly, similar to CLIC1^17,23,25^, EGFRvIII impacts various aspects of tumourigenesis not only in the cell of origin, but in the neighbouring cells, also shown to be mediated via EVs.^26^ Thus, we first questioned whether CLIC1 is required for the horizontal propagation of EGFRvIII to the neighboring cells. By extracting EVs from CTL BTSCs and their corresponding CLIC1 CRISPR cells, we evaluated EGFRvIII protein expression levels in the cell lysates and EV fractions. Immunoblotting analysis revealed that EGFRvIII levels were significantly reduced in CLIC-deleted EVs, suggesting a role for CLIC1 in mediating the horizontal propagation of EGFRvIII to the neighboring cells (**Fig. 6a-c**, **Supp. Fig. 8)**. Next, we set out to investigate the electrophysiological properties of tmCLIC1 in three distinct human BTSC lines including BTSC73, BTSC147, and BTSC30. To assess basal level tmCLIC1 levels, we used IAA94, a known blocker of CLIC1. The tmCLIC1 current was measured as the IAA94-sensitive current, defined as the difference between the initial current and the current after drug perfusion. We conducted an electrophysiological evaluation of tmCLIC1 levels at +40mV on the membranes of CLIC1 CRISPR and control BTSCs. Our results revealed that the tmCLIC1 currents were significantly reduced in the CLIC1 CRISPR lines compared to the CTL BTSCs (**Fig. 6d**). To assess whether OSMR is required for the tmCLIC1 mediated ionic conductance, we induced KD of OSMR using short hairpin RNA (shOSMR) in different cell lines and subjected the OSMR knockdown (KD) and control (CTL) Scrambled (SCR) cells to patch clamp experiments to evaluate whether reduced OSMR expression levels alter Cl^−^ currents. Cell currents were assessed both prior to (baseline) and following the addition of 100 mM of CLIC1 inhibitor, IAA94 to the bath solution (**Fig. 6e-j**). The tmCLIC1 current was measured as the IAA94-sensitive current, defined as the difference between the initial current and the current after drug perfusion. The corresponding current-voltage relationship was recorded in each group. Strikingly, our data revealed that inhibition of OSMR significantly attenuated the tmCLIC1-associated Cl^−^ current (**Fig. 6i-k).** Our data reveals a bidirectional OSMR-CLIC1 cross talk in regulation of CLIC1-mediated ionic conductance in BTSCs, leading to the question of whether tmCLIC1 impacts EGFRvIII/STAT3 signalling. To address this question, we generated and employed a monoclonal CLIC1 antibody (tmCLIC1omab^®^) that specifically inhibits tmCLIC1 and its associated ionic currents (**Fig. 6l-p**, **supp. Fig. 9a-d**). We first assessed the tmCLIC1 antibody in tumour assays in which BTSCs xenografted in the flanks of SICD mice were treated with tmCLIC1 antibody or IgG control. Strikingly, tmCLIC1 significantly suppressed tumour growth in the antibody treated group (**Fig. 6q-s**). Next, we treated different glioma cells and BTSCs with the tmCLIC1omab or IgG control, followed by an analysis of EGFRvIII and STAT3 phosphorylation in response to the ligand OSM. Our data revealed that both the EGFR phosphorylation at tyrosine (Y) 1068 and STAT3 phosphorylation at Y705 were significantly attenuated upon inhibition of the CLIC1-mediated Cl-current by a tmCLIC1 antibody in EGFRvIII expressing BTSC 147 and 73. Importantly, tmCLIC1omab^®^ had no impact on EGFRvIII-STAT3 signalling in BTSC30 that naturally does not harbour the EGFRvIII or elevated STAT3 expression (**Fig. 6t-y, Supp. Fig. 10**).

Taken together, we provide data that tmCLIC1 regulates OSM/OSMR-STAT3 and OSMR/EGFRvIII oncogenic signalling and at the same time OSMR promotes tmCLIC1 function. We establish that the cooperation of OSMR with tmCLIC1 is required to maintain key oncogenic pathways in GB.

### CLIC1 drives GB tumourigenesis and maintains EGFRvIII/EGFR phosphorylation *in vivo*

Our findings on a central role of CLIC1 in regulation of OSMR/EGFRvIII interaction and EGFRvIII and STAT3 phosphorylation led us next to examine whether CLIC1 contributes to GB tumourigenesis *in vivo*. We conducted intracranial tumour assay using BTSC73, BTSC147, and corresponding CLIC1 CRISPR BTSCs lines. Our data revealed that deletion of CLIC1 significantly impaired tumourigenesis, and prolonged survival (**Fig. 7a-h**). We next subjected tumour sections obtained from the control mice as well as the smaller tumours from the CLIC1 CRISPR group to H&E staining (**Fig. 7i,j**) and immunostaining analysis using a phospho-Y1068-EGFR antibody that can detect both the phosphorylated wild type (WT) EGFR and EGFRvIII (**Fig. 7k**). Strikingly, we found that the phosphorylation of EGFR/EGFRvIII was robustly attenuated in the mice group xenografted with CLIC1 CRISPR BTSCs. Independently, we subjected different CLIC1 CRISPR and control BTSCs to immunoblotting experiments and analyzed the phosphorylation events using antibodies to STAT3-Y-705 and EGFRvIII. Strikingly, similar to results obtained with tmCLIC1moab, we established that genetic deletion of CLIC1 significantly impairs the phosphorylation of STAT3 and EGFRvIII. (**Fig. 7l-s**). Our data suggest that CLIC1 tightly maintains the activation of EGFRvIII and STAT3.

**Figure 7.**
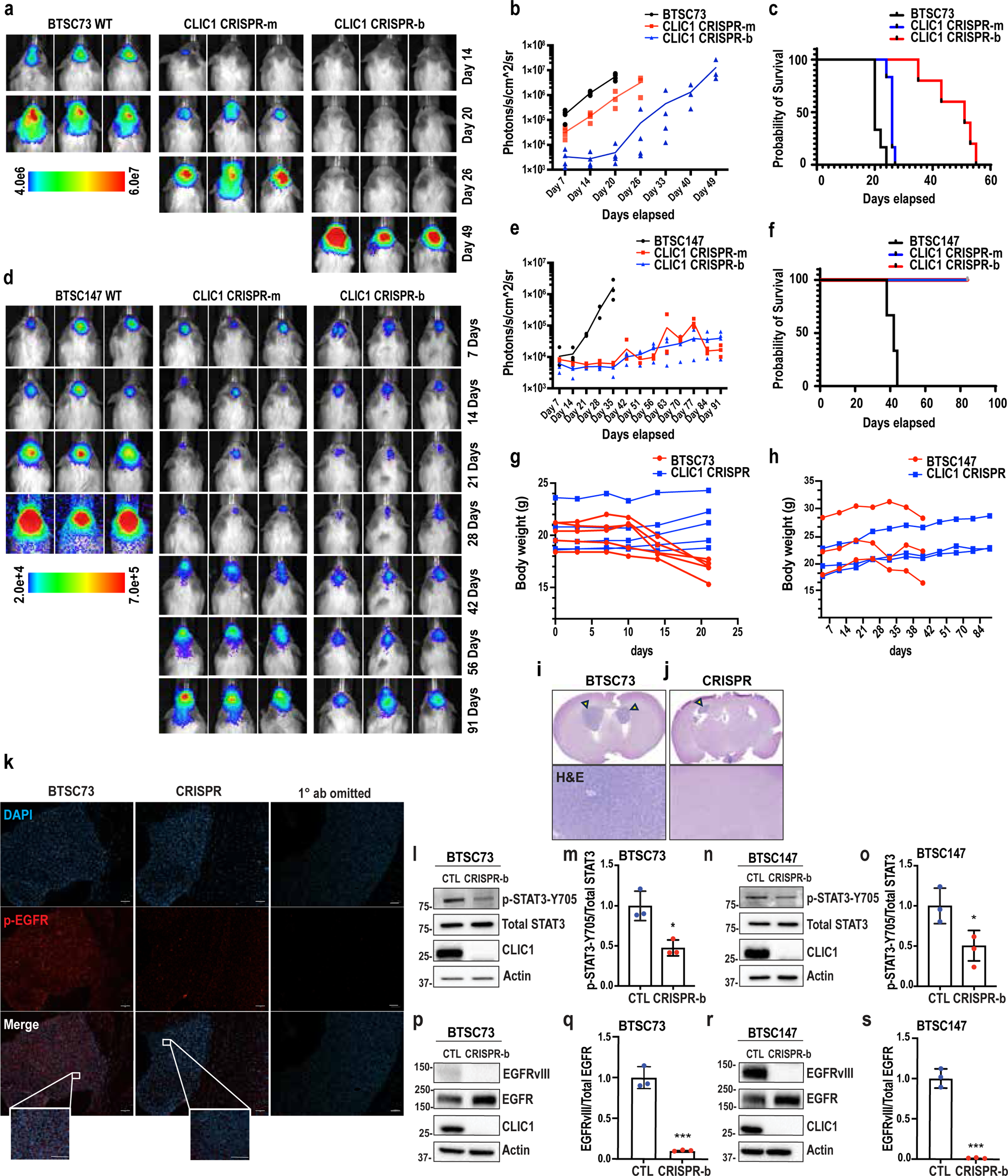
Genetic deletion of CLIC1 impairs tumourigenesis, expands lifespan and suppresses EGFR/EGFRvIII activation. **a-h,** CLIC1 CRISPR BTSCs (#73,) control BTSC73, CLIC1 CRISPR BTSCs (#147), and control BTSC147 were xenografted intracranially into 8 weeks old SCID male mice. IVIS imaging was conducted to evaluate tumour volume (**a,d**) and signal was quantified (**b,e**). Kaplan-Meier survival plots were graphed to evaluate animal lifespan in each group (**c,f**). Log-rank test (BTSC73) (*P* < 0.0001, *n* ≥ 5 mice), BTSC#73 vs. CLIC1 CRISPR-a (*P* = 0.0011), BTSC#73 vs. CLIC1 CRISPR-b (*P* = 0.001), CLIC1 CRISPR-a vs. CLIC1 CRISPR-b (*P* = 0.0014). Log rank test for BTSC#147, (*P* = 0.0078, n=3), BTSC#147 vs. CLIC1 CRISPR-a (*P* = 0.0246), BTSC#147 vs CLIC1 CRISPR-b (*P* = 0.0246), CLIC1 CRISPR-a vs CLIC1 CRISPR-b (non-significant). Representative panels for changes in body weight following BTSC implantation are shown (**g-h**). **i-k**, Brain sections depicting tumour in control BTSC73 and CLIC1 CRISPR BTSC73 were subjected to H&E staining (**i,j**) or immunostaining with a p-EGFR-Y1068 antibody (**k**). **l-s**, CLIC1 CRISPR and control BTSCs were subjected to immunoblotting using antibodies to P-STAT3-Y705, total STAT3, EGFRvIII, and total EGFR. ACTIN was used as loading control. Densitometric quantification of p-STAT3 to total STAT3 (**m,o**) and EGFRvIII to total EGFR (**q,s**) are presented. Statistical analysis was performed using a student t-test. n=3 biological replicates. BTSC73 (**m**): \**P* = 0.0121; BTSC147 (**o**): \**P* = 0.0423; BTSC73 (**q**): \*\*\**P* = 0.0003; BTSC147 (**s**): \*\*\**P* = 0.0001 for each pairwise comparison.

## Discussion

In this study, we comprehensively mapped the OSMR interactome to identify its high confidence binding partners under different conditions leading to a better understanding of OSMR’s multifaceted roles in GB progression. Through systematic protein-protein interaction mapping using MaMTH-HTS, followed by counter-screening and loss- and gain-of-function studies, we demonstrate that tmCLIC1 interacts with OSMR physically, independently of the genetic background of the cells, and acts as a critical regulator of diverse oncogenic pathways driven by OSMR, EGFRvIII, and STAT3. We demonstrate a bidirectional relationship for tmCLIC1-OSMR interaction, which plays a pivotal role in regulating ion conductance, thereby driving GB progression.

CLIC1 has been implicated in multiple malignancies, including lung cancer^27^, pancreatic adenocarcinoma^28^, epithelial ovarian cancer^29^, and medulloblastoma.^30^ It is associated with poor prognosis in GB and is abundantly expressed in CSCs.^17,31^ Previous studies have highlighted the role of tmCLIC1 in cellular proliferation and viability, with genetic deletion leading to cell swelling, mitotic defects, and reduced proliferation in medulloblastoma cells.^30^ tmCLIC1 may act as a cell cycle accelerator by mediating cell volume changes. For instance, during the prophase to metaphase transition, cells undergo a significant volume decrease, reaching a minimum size at metaphase. This reduced volume is preferred by the cell and is referred to as pre-mitotic condensation.^32,33^ Pre-mitotic condensation requires the efflux of Cl-, which may be mediated by tmCLIC1. tmCLIC1 has also been shown to play a prominent role in ROS production^18,34,35^, and cell cycle regulation.^36^ For example, IAA94-mediated inhibition of tmCLIC1 is shown to prolong the G1 phase, thereby extending the overall cell cycle duration.^37^ In agreement with these findings, our data show that CLIC1 deletion suppresses BTSC self-renewal and proliferation, suggesting its role in cell cycle regulation. Given tmCLIC1’s established function in maintaining redox balance^38^, its knockdown likely elevates oxidative stress and disrupts redox homeostasis, thereby impairing BTSC proliferative capacity and self-renewal potential. These findings reinforce the idea that CLIC1 is a critical regulator of GB cell division and tumour aggressiveness. The interplay between ROS, tmCLIC1-generated chloride currents, combined with evidence of tmCLIC1/OSMR interaction, highlights the need to investigate CLIC1’s role in CSCs’ metabolism in the context of GB. Whether similar to OSMR^5^, tmCLIC1 regulates oxidative phosphorylation (OXPHOS) to maintain BTSCs and buffer ROS in the mitochondria remains to be investigated.

We further demonstrate a crucial role for CLIC1 in maintaining EGFRvIII intracellularly. This effect may also be attributed to CLIC1’s role in tumour microenvironment modulation, as it is positively associated with stromal activation, epithelial-mesenchymal transition (EMT), and angiogenesis, hallmarks of GB progression.^39^ Chloride channels, including CLIC1, have been implicated in innate and adaptive immune responses, particularly in phagocyte-mediated immunity. CLIC1 is required for phagosomal acidification and antigen processing, which are crucial for CD4⁺ T cell activation.^39–41^ CLIC1-mediated regulation of macrophage-driven inflammation and ROS production may facilitate the establishment of a tumour-promoting microenvironment conducive to tumour growth and metastasis. Furthermore, CLIC1 plays a role in the innate immune system through its involvement in the activation of NLRP3 inflammasome.^42^ Aberrant activity of the NLRP3 inflammasome has been implicated in the progression of several cancers including glioma.^43,44^ Additionally, tmCLIC1 modulates Cl⁻ currents in activated microglia, particularly in response to β-amyloid protein (Aβ) stimulation.^35^ Interestingly, OSMR and its ligand OSM operate in a paracrine fashion in which OSM secreted via immune cells typically binds the receptor on the tumour cells to direct oncogenic signalling. These findings suggest that OSMR-CLIC1 cross talk may influence immune surveillance, an area that requires future investigation.

It is also possible that loss of CLIC1 induces a cell shift towards a more differentiated phenotype. Previous research on cell fate dynamics supports this notion. Stockhausen et al. demonstrated that induced differentiation of BTSCs with serum leads to loss of EGFRvIII expression and decreased tumourigenic potential.^45^ In another study by Gritti et al., CLIC1 expression was lost upon induction of differentiation with FBS.^46^ These results suggest that both EGFRvIII and CLIC1 are important in maintaining stemness. Similarly, in another study by He et al., the chloride channel, Ano1, was found to limit the differentiation of epithelial progenitors towards cells of the secretory lineage.^47^ Further studies are required to determine if the interaction of CLIC1 with EGFRvIII aids in establishing a cellular network that suppresses the BTSCs differentiation.

In this study, we provide data showing that CLIC1 regulates the horizontal transfer of EGFRvIII via EV. We observed a striking attenuation of EGFRvIII in EVs derived from CLIC1 CRISPR cells, suggesting that CLIC1 may regulate EGFRvIII packaging, thereby influencing neighboring cells. In support of this model, tmCLIC1 vesicle transfer has been implicated in glioma stem cell proliferation.^23^ Furthermore, blocking of the chloride current in stem-like cells through the re-purposing of known pharmacological drugs, suggests that tmCLIC1 could be a promising target for therapy.^46,48^ Similarly, tumour cells in GB are shown to impact neighboring cells via transferring EGFRvIII through EV.^26^ Another study in gastric cancer revealed that the presence of CLIC1 in EVs is associated with resistance to vincristine by transferring resistance traits to neighboring cells.^24^ Whether tmCLIC1 functional expression represents an adaptive mechanism enabling drug resistance in GB remains an exciting area of research to be explored.

Our findings have established that tmCLIC1 interaction with OSMR and EGFRvIII is essential for sustaining oncogenic signaling. Using a monoclonal CLIC1 antibody that can only target tmCLIC1, we establish the significance of tmCLIC1, however, the functional contribution of cytoplasmic CLIC1 and its involvement in mitochondrial and metabolic pathways remains to be investigated in GB.

In conclusion, our findings establish tmCLIC1 as a central player in GB pathogenesis, orchestrating cytokine networks and enhancing the RTK signalling. Its interaction with OSMR and EGFRvIII is critical for sustaining BTSC self-renewal and GB tumourigenesis. These results highlight tmCLIC1 as a promising therapeutic target, suggesting that its inhibition, in combination with standard GB treatments, may lead to improved patient outcomes. Importantly, our data suggests that inhibiting the interaction of CLIC1 and OSMR in GB using small inhibitory peptides serves as a promising therapeutic avenue for GB.

## Materials and methods

### Brain tumour Stem Cell (BTSC) and generation of transgenic cell lines

The human BTSC lines 73, 147, 12, and 30 were generously provided by Dr. Samuel Weiss at the University of Calgary. They were generated in accordance with the University regulation from excess or discarded tissue collected during surgery from adult patients following informed consent from patients. BTSCs were characterised for major mutations (**Supp. Table 2**) and were cultured and maintained, as described in Supplementary Materials and methods section. Transgenic CRISPR BTSCs were generated, using methods previously described.^49^ To design the gRNA, Off-Spotter software, version 0.2.2 (https://cm.jefferson.edu/Off-Spotter/), was used. Two guide RNA strands (forward and reverse complement) were generated to target exons 5-9 of the CLIC1 gene. To generate the construct, the Golden Gate Assembly Cloning strategy^50^ was used in which gRNA1 and gRNA2 were cloned into pL-CRISPR.EFS.GFP (Addgene plasmid #57818), and pL-CRISPR.EFS.tRFP (Addgene plasmid, #57819), plasmids, respectively. Plasmids were sequenced (Genome Quebec), verified and electroporated [1300 volts by the AMAXA nucleofector 2b device (Lonza, #AAB1001) to deliver 3 µg of each plasmid construct (gRNA1-GFP and gRNA2-RFP) into ∼2 million BTSC147 or BTSC73]. Electroporated cells were cultured in a T-75 low attachment flask at 37°C in a 5% CO_2_. 48 hours after electroporation, BTSC spheres were dissociated into a single cell suspension using Accumax dissociation solution (Innovative Cell Technologies, #AM105) and subjected to Fluorescent Activated Cell Sorting (FACS) analysis using the BD FACSAriaTM Fusion (BD Biosciences) to sort for double positive GFP and RFP cells. Sorted cells were plated at a density of 1 cell/well into two 96-well plates containing 100 µl of BTSC media. Wells were monitored every two days to assess sphere formation and clonal samples were collected from multiple positive clones and were subject to genomic DNA isolation. Isolated DNA was analysed by PCR using internal primers and external primers to the gRNA-guided CAS9 cut site, designed using Primer3Plus software, version 3.3.0 (https://www.primer3plus.com/), to determine monoallelic, biallelic, or non-deletion control clones. Monoallelic deletion clones in both BTSC73 and BTSC147 cell lines were determined by PCR in which the presence of one internal and one external band on a 2% Agarose gel, was observed, while a biallelic deletion was determined by a single external band. Knockout or knockdown of CLIC1 was validated using RT-qPCR and WB to assess gene and protein expression. Since there we no clones identified that presented no cuts in the CLIC1 gene following incubation with the CRISPR-CAS9 construct, which could have been used as a control for CAS9, the chosen control for experiments using the CRISPR-CAS9-CLIC1 cells was that of the parental BTSC73 or BTSC147 line. The gRNA and primer sequences used in this experiment are listed in **Supp. Table 4**. Finally, short Interfering RNA (siRNA) was used to generate transient knockdown (KD) in patient-derived BTSCs. BTSCs were processed into a single-cell suspension. ON TARGET-plus SMART pool human CLIC1 siRNA (Dharmacon, #L-009530-00-0005) at a concentration of 100 nM, and ON TARGET-plus non-targeting pool (Dharmacon, #D-001810-10-05), were employed, as described in Supplemental Materials and methods section.

### Mammalian Membrane Two-Hybrid-High Throughput Screen (MaMTH-HTS)

To identify binding partners of OSMR in the presence and absence of EGFRvIII, Mammalian Membrane Two-Hybrid (MaMTH) High Throughput Screen (HTS) technology was used as described.^16,51^ Briefly, plasmid expressing OSMR ‘Bait’ protein with C-terminally fused MaMTH-HTS Bait tag (Cub-GAL4TF-P2A-tagBFP) alone or alongside plasmid expressing EGFRvIII (fused with 3xFLAG at its N-terminus) was transfected into a pooled ‘Prey’ library of HEK293T MaMTH-HTS reporter cell lines. The library of reporter cell lines stably expressed members of the Human ORFeome V8.1 collection (∼8000 open reading frames - ORF’s) fused to MaMTH-HTS Prey tag (Nub) at their N-terminus and P2A-mCherry at their C-Terminus and contained chromosomally integrated GFP reporter under the control of the GAL4 transcription factor (GAL4TF) promoter. Transfections were performed using X-tremeGene™ 9 Transfection Reagent (Roche, XTG9-RO) as specified by the manufacturer protocol. In order to induce Bait (**Supp. Fig. 1**) and Prey expression, cells were grown for 2-3 days in the presence of 0.5 µg/ml Tetracycline in the following conditions: 37°C, 5% CO_2_ in DMEM containing 10% FBS, and 1% Penicillin/Streptomycin media. Cells were then harvested by trypsinization and resuspended at a concentration of 1-2 × 10^6^ cell/ml in Basic Sorting Buffer (1X PBS, 5 mM EDTA, 25 mM HEPES pH 7.0, 1% BSA) and subjected to sorting by Flow Cytometry using BD FACSMelody (BD Biosciences). Cells were sequentially selected according to the following: tag-BFP fluorescence indicating ‘Bait’ expression, mCherry fluorescence indicating ‘Prey’ expression, and GFP fluorescence indicating Bait-Prey interaction. Cells were collected for analysis in DMEM containing 25% FBS and centrifuged pellets were processed in Phire Tissue Direct Dilution Buffer (ThermoFisher Scientific). Amplification of ORFs was done using Phire Tissue Direct PCR Master Mix (ThermoFisher Scientific) and products were purified using QIAquick PCR Purification Kit (Qiagen). Purified PCR products were subjected to Nextera XT library preparation and deep sequencing using the Illumina HiSeq 2500 system (150 bp single read). Sequencing data was processed and hits were identified using custom software developed using R programming language and integrated Bowtie2 alignment tool, version 2.5.2.^52^

### Counter-screen assay

BTSCs were transfected with siRNA using RNAiMax in a 96-well plate. Briefly, 1 pmol of siRNA and 0.1 µl of RNAiMax were diluted in 5 µl of Opti-MEM and incubated separately for 5 minutes. The solutions were then gently combined and incubated at room temperature for 15 minutes to form transfection complexes. A total of 1,000 BTSCs per well were seeded in 80 µl of NeuroCult medium, followed by the addition of 10 µl of the siRNA-RNAiMax complex with gentle shaking. After incubation at 37°C for 5 hours, 90 µl of NeuroCult medium was added to each well, and cells were further incubated at 37°C for 48 hours. Transfected cells were then subjected to the PrestoBlue assay to assess cell viability.

### Duolink Proximity Ligation Assay (PLA)

Proximity Ligation Assay was conducted following the manufacturer’s protocol using the Duolink *In Situ* Red Starter Kit (Sigma, #DUO92101) as optimized by our group^53^, and described in the Supplemental Materials and methods section.

### Extreme Limiting Dilution Assay (ELDA)

ELDA was performed as previously described^54^, with detailed methods presented in the Supplemental Materials and methods section.

### 5-ethynyl-2’-deoxyuridine (EdU) Assay

Cell proliferation of BTSC73, BTSC147, and their related CLIC1 CRISPR KD cell were assessed using the Click-iT™ Plus EdU Alexa Fluor™ 488 Flow Cytometry Assay Kit (Invitrogen, Cat #C10632). According to the manufacturer’s protocol, cells were incubated with 10μM EdU for 2 hours at 37 °C to label newly synthesized DNA. Following this step, cells were fixed, permeabilized, and washed twice with a permeabilization and wash buffer provided in the kit. The incorporated EdU was then detected with the Alexa Fluor 488 azide via a copper-catalyzed click reaction. Following a final two washes, samples were analyzed using Cytek Aurora Flow Cytometry System (5) Lasers/ (64) Fluorescence Channels/ (67) Detector (Cytek Biosciences# 62352, USA). Data were analyzed using FlowJo™ v10 Software.

### In vitro binding assay

HEK293T cells were seeded in 6-well plates at approximately 60–70% confluency one day prior to transfection. Cells were transfected using PEI with plasmids encoding Myc-tagged truncated CLIC1 constructs, CLIC1(1–58) and CLIC1(1–178), together with expression vectors for OSMR and/or EGFRvIII. The total amount of DNA per well was normalized with empty vector to ensure equal transfection conditions across samples. Twenty-four hours post-transfection, cells were washed twice with ice-cold phosphate-buffered saline (PBS) and lysed on ice in lysis buffer (20 mM Tris-HCl, pH 7.5, 150 mM NaCl, 0.5% NP-40, 1 mM EDTA) supplemented with protease and phosphatase inhibitor cocktails. Lysates were incubated on ice for 20 minutes and clarified by centrifugation at 12,000 × g for 15 minutes at 4°C. The supernatants were collected for immunoprecipitation (IP). 500 µg of total cell lysate was incubated with 1 µg of anti-Myc antibody overnight at 4°C with gentle rotation to IP Myc-tagged CLIC1 and associated proteins. The following day, Protein G Dynbeads were added and incubated for an additional 2 hours at 4°C. Beads were then washed three times with lysis buffer. IPed proteins were eluted by boiling the beads in 2x SDS sample buffer for 5 minutes and subjected to SDS–PAGE, followed by Western blotting (WB). Proteins were transferred to PVDF membranes, which were blocked in 5% BSA in TBST (20 mM Tris-HCl, 150 mM NaCl, 0.1% Tween-20, pH 7.4) for 1 hour at room temperature. Membranes were then incubated overnight at 4°C with primary antibodies against Myc (for CLIC1), OSMR, and EGFRvIII, followed by HRP-conjugated secondary antibodies for 2 hours at room temperature, and were visualized using ECL. A fraction of total lysate (5% of total protein used for IP) was analyzed in parallel to confirm expression of transfected constructs (input control).

### Stereotaxic injections and bioluminescent imaging

Stereotaxic surgeries were conducted by injecting ∼3 × 10^5^ luciferase expressing CLIC1 CRISPR BTSCs and control BTSCs into the right striata (0.8 mm lateral to the bregma, 1 mm dorsal, and 2.5 mm from the pial surface). Prior to injection, BTSCs were dissociated into single-cell suspensions in serum-free, antibiotic-free medium and tested for luciferase activity using IVIS imaging system. Kaplan-Meier survival plots were generated by collecting mice at the point of reaching the end stage (major body weight loss, dehydration, hunched back, piloerection, and lethargy). Median survivals were calculated using a log-rank test with GraphPad Prism, following previously described methods.^55^ Briefly, the Kaplan-Meier approach estimated the probability of survival at each time point. Before any time elapses, all mice were considered “at risk”, however, no deaths occurred. Therefore, the probability of survival was input as a value of 1. Subsequently, information about the next elapsed time (day 7) included information about the how many participants experienced the event of interest (death). From this information, a survival probability was calculated using the formula: S_t+1_ = S_t_*((N_t+1_-D_t+1_)/N_t+1_), where S_t_ represents survival probability, N_t_ represents number at risk, D_t_ represents number of events. The survival probability for each subsequent time point is calculated, and a stair-step survival curve is plotted. For tumour volume assessment, the mice received intraperitoneal injections of 200 µl of 15 mg/ml d-luciferin (ThermoFisher Scientific, #88292), underwent anesthesia via isoflurane inhalation, and were then subjected to weekly bioluminescence imaging using a CCD camera (IVIS, Xenogen). Subsequent collection and analysis of all bioluminescent data were performed utilizing Living Image 2.0 software (PerkinElmer, MA, USA). Briefly, after initialization of the IVIS Spectrum system and acquiring an image, the images were analyzed using “ROI tools” in the tool palette, selecting the area of interest. The IVIS imager captures pixels with specific photon intensity values, with brighter areas corresponding to higher photon detection, indicative of tumour induction.

### Design and generation of the tmCLIC1 monoclonal antibody

We designed and generated tmCLIC1 mAb to specifically target the NH₂-terminal region of the CLIC1 protein. Polyclonal antibodies were initially generated through a 138-day immunization protocol in which mice were immunized with a synthetic peptide corresponding to the NH₂-terminus of CLIC1 (NH₂-EQPQVELFVKAGSDGAKIGNC-COOH), conjugated to ovalbumin (OVA) and dissolved in a buffer composed of 0.1 M glycine (pH 3) and 1 M Tris-HCl (pH 8). Antibody specificity was confirmed by the supplier using an ELISA assay. Hybridoma cells producing antibodies were sorted as single cells to isolate monoclonal clones. A total of 44 distinct monoclonal antibodies, derived from all four immunized mice, were sent to our laboratory for further screening. Among these, we selected the clone showing the highest specificity and sensitivity. Sequencing of the selected clone revealed strong binding affinity to the QVELF motif of the immunizing peptide, a sequence conserved across several species and uniquely expressed by tmCLIC1, but not by other members of the CLIC protein family. Multiple experiments were conducted to assess the specificity and functional effectiveness of tmCLIC1omab in inhibiting tmCLIC1 activity (**Fig. 6l-n, supp. Fig. 9a,b**). We also demonstrated that the antibody retains its effect on primary human glioblastoma cells even after short incubation periods (**Fig. 6o**), and that it is efficiently internalized into cells, as shown using the pHrodo™ dye assay (**supp. Fig. 9c,d**). Additionally, we evaluated the *in vivo* tolerability of tmCLIC1omab in a mouse model of human colorectal cancer. Mice received biweekly injections of the antibody for three weeks. Body weight monitoring showed no significant difference between tmCLIC1omab-treated mice and those receiving an isotype control, indicating that the antibody is well tolerated *in vivo* (**Fig. 6p**) (The expanded version can be found in the Supplemental Materials and methods section).

### Patch-Clamp Experiment

Patch electrodes (BB150F-8P with filaments, Science Products), with a diameter of 1.5 mm, were pulled from hard borosilicate glass on a Brown-Flaming P-87 puller (Sutter Instrument, Novato, CA) and fire polished to a tip diameter of 1-1.5 µm and an electrical resistance of 3-4 Mꭥ. Cells were voltage-clamped using an Axopatch 200B amplifier (Axon Intrument) in whole-cell configuration in which After formation of giga-seal, the membrane patch is disrupted providing a direct low resistance access to cell interior, allowing recordings from ion channels of whole cell. The voltage step protocol used to isolate current/voltage relationships consisted of 800 ms pulses from -60 mV to +60 mV (20 mV voltage steps). The holding potential was set according to the resting potential of the single cell (between -40 and -80 mV). tmCLIC1-mediated chloride currents were isolated from other ionic currents by perfusing IAA94 (100 μM) dissolved in the bath solution and by mathematical subtraction of the residual current from the control. Solution used are the following: bath solution (mM): 125 NaCl, 5.5 KCl, 24 HEPES, 1 MgCl2, 0.5 CaCl2, 5 D-Glucose, 10 NaOH; pH 7.4; Pipette solution (mM): 135 KCl, 10 Hepes, 10 NaCl; pH 7.4. Analysis was performed using Clampfit 10.2 (Molecular Devices) and OriginPro 9.1. tmCLIC1-mediated current (IAA94-sensitive current) was measured by analytical subtraction of residual ionic current after addition of inhibitor from total current (I_TOT_) of the cell at each membrane potential tested. Current/voltage relationship were constructed plotting the averaged current density of the least 100 ms of the pulse against the corresponding membrane potential. Current density (pA/pF) results from the ratio between the ionic current (pA) and cells capacitance (pF). Statistical analyses were performed comparing the slopes (proportional to channel conductance) of the I/V curves of the different groups.

### Generation of monoclonal tmCLIC1 antibody

The tmCLIC1omab^®^ antibody was designed (license# 11665M), targeting the N-terminal of the CLIC1 protein derived from mice. Polyclonal antibodies were obtained through a 138-day immunization protocol in which mice are immunized against a NH_2_-CLIC1 synthetic peptide conjugated to OVA (ovalbumin) (NH_2_-EQPQVELFVKAGSDGAKIGNC-COOH) (Glycine 0.1 M pH 3, Tris-HCl 1 M pH 8). Hybridoma cells that produced antibodies were sorted in single cells to obtain monoclonal antibodies. tmCLIC1omab^®^ was used at a final concentration of 3.5 µg/ml.

### Extracellular vesicle isolation and validation

To sequentially isolate EVs, the cell supernatant was centrifuged at 200 x g for 10 minutes, 300 x g for 10 minutes, 2,000 x g for 10 minutes, 10,000 x g for 30 minutes, and 100,000 x g for 2 hours, respectively (Optima XPN-100 ultracentrifuge, Beckman Coulter, USA). A beige/white pellet of EVs and a clear supernatant were evident post-centrifugation. After removing the supernatant and resuspending the small EV pellet in 1 ml PBS, the resulting pellet was then washed and centrifuged at 49,000 rpm for 30 minutes (Optima MAX 130K Refrigerated Benchtop Ultracentrifuge, Beckman Coulter, USA). Isolated EVs were then visualized, measured, counted, and characterized in the range of 10-2,000 nm using a Nanoparticle Tracking Analyzer (NTA) (Particle Metrix ZetaView® Nanoparticle Tracking Analysis).^56,57^ We used size-mode analysis with ZetaView software (version 8.02.28) following calibration with polystyrene beads (105 and 500 nm). Samples were analyzed at a minimum of 9 camera positions with 2-second video length at 21°C.

### Bioinformatic analysis

Novel interactions from MAMTH screen were integrated with the known interactions from the Integrated Interaction Database IID ver. 2021-05^58^ and further annotated with Gene Ontology Biological Process and Cellular Component (release 2022-11-03), and a subset of pathways from the PathDIP database ver. 4.^59^ Resulting network was built, annotated and analyzed using NAViGaTOR ver. 3.0.19.^60^

### Statistical analysis

Statistical analysis between conditions was performed using a Student *t*-test or one-way ANOVA in which the mean of n=3 replicates were compared between two conditions to assess significance. Analysis was undertaken with the aid of GraphPad software 7. Data is shown as mean with standard deviation (mean ± SD). p-values less than 0.05 were considered significant and were marked with an asterisk as follows, \**P* < 0.05; \*\**P* < 0.01; \*\*\**P* < 0.001. Precise p values are provided in the figure legends, unless noted otherwise.

## Acknowledgements

This work was supported by Canadian Institute for Health Research (CIHR) grants (# 157480, #157797), Canada Foundation for Innovation (CFI) (#156794), Ontario Research Fund (ORF) (#156796), Canada Research chair (CRC) (#156787, 156789), to A.J.-A, and CIHR grants to AJA, I.S., J.R. (# 162198, #519474) and a AIRC IG grant (#24758) to M.M.. A.J.-A. is a Canada Research Chair at the University of Ottawa and is supported by CIHR, Natural Sciences Research Council (NSERC), Cancer Research Society (CRS). I.S. is supported by CFI (#225404, #30865), ORF (RDI #34876, RE010-020), and (NSERC RGPIN-2024-04314). IJ was supported in part by funding from NSERC (RGPIN-2024-04314), CIHR (#519474), CFI #225404, #30865, and ORF (RDI #34876, RE010-020). HS is supported by NCATS (R03TR005313), NHLBI (R01HL133050). The funders had no role in study design, data collection and analysis, decision to publish, or preparation of the manuscript.

