## Supplemental file for "An OSMR-CLIC1 cross talk drives key oncogenic pathways in glioblastoma"

**Supplementary Information:**

**1. Supplementary Tables 1 to 5**

**- Supplementary Table. 1**

**- Supplementary Table. 2**

**- Supplementary Table. 3**

**- Supplementary Table. 4**

**- Supplementary Table. 5**

**2. Supplementary Figures 1 to 10**

**- Supplementary Fig. 1.**

**- Supplementary Fig. 2**

**- Supplementary Fig. 3**

**- Supplementary Fig. 4**

**- Supplementary Fig. 5**

**- Supplementary Fig. 6**

**- Supplementary Fig. 7**

**- Supplementary Fig. 8**

**- Supplementary Fig. 9**

**- Supplementary Fig. 10**

**3. Supplementary Material and methods**

**4. References**

**Supplementary Table. 1. Common candidate binding partners to OSMR-unique and OSMR/EGFRvIII-unique**. High confidence binding partners to OSMR regardless of cell's genetic background was extracted for follow up analysis. The cellular localization of the common candidates is illustrated in the last column.

| # | Common candidate binding partners | Detected by MaMTH | Novel MaMTH PPI | localization |
|---|---|---|---|---|
| 1 | Argonaute 3 (AGO3) | high conf | y | cytoplasm |
| 2 | B-cell receptor-associated protein 31 (BCAP31) | high conf | y | plasma membrane |
| 3 | BCL2 Like 14 (BCL2L14) | high conf | y | cytoplasm |
| 4 | Chromosome 8 open reading frame 44 (C8orf44) | high conf | y | nucleus |
| 5 | C-type lectin domain family 2 member B (CLEC2B) | high conf | y | plasma membrane |
| 6 | Chloride Intracellular Channel 1 (CLIC1) | high conf | y | plasma membrane |
| 7 | Cyclic AMP-responsive element-binding protein 3 (CREB3) | high conf | y | Golgi apparatus |
| 8 | Chondroitin sulfate N-acetylgalactosaminyltransferase 2 (CSGALNACT2) | high conf | y | Golgi apparatus |
| 9 | Chromosome X open reading frame 1 (CXorf1) | high conf | y | cytoplasm |
| 10 | Decorin (DCN) | high conf | y | Golgi apparatus |
| 11 | Guanylate cyclase (GC) | high conf | y | cytoplasm |
| 12 | Glutaredoxin (GLRX) | high conf | y | nucleus |
| 13 | GC-rich promoter binding protein 1 (GPBP1) | high conf | y | plasma membrane |
| 14 | Histone H3, clustered gene 1 (H3C1) | high conf | y | nucleus |
| 15 | Inner Mitochondrial Membrane Peptidase Subunit 2 (IMMP2L) | high conf | y | mitochondrion |
| 16 | Leucine Rich Repeats and IQ Motif Containing 1 (LRRIQ1) | high conf | y | Cytoplasm and plasma membrane |
| 17 | Microtubule-Associated Protein 4 (MAP4) | high conf | y | plasma membrane |
| 18 | N-acetyltransferase 8 (NAT8) | high conf | y | endoplasmic reticulum |
| 19 | Nurim (NRM) | high conf | y | nucleus |
| 20 | phosphatidylinositol glycan anchor biosynthesis class K (PIGK) | high conf | y | endoplasmic reticulum |
| 21 | Psoriasis susceptibility 1 candidate 1 (PSORS1C1) | high conf | y | |
| 22 | Phosphatidylserine Synthase 1 (PTDSS1) | high conf | y | endoplasmic reticulum |
| 23 | Ras-related protein Rab-1B (RAB1B) | high conf | y | Golgi apparatus |
| 24 | SET domain-containing 7 histone lysine methyltransferase (SETD7) | high conf | y | nucleus |
| 25 | Solute Carrier Family 35 Member B3 (SLC35B3) | high conf | y | Golgi apparatus |
| 26 | Small Integral Membrane Protein 21 (SMIM21) | high conf | y | plasma membrane |
| 27 | ST3 beta-galactoside alpha-2,3-sialyltransferase 4 (ST3GAL4) | high conf | y | Golgi apparatus |
| 28 | TRAF family member associated NFKB activator (TANK) | high conf | y | cytoplasm |
| 29 | Transmembrane protease, serine 6 (TMPRSS6) | high conf | y | plasma membrane |
| 30 | Zona Pellucida Glycoprotein 1 (ZP1) | high conf | y | plasma membrane |

**Supplementary Table. 2. Patient derived BTSCs and their genetic signature**. Four different patients derived BTSCs were employed in this study. The gender and age of the glioblastoma patients and their characterization for key genetic mutations are shown. M, male; wt, wild type; mut, mutant; vIII, EGFR variant III; homo del, homozygous deletion; NA – not available.

| Cell line | Sex | Age | *EGFR* | *TP53* | *PTEN* | *IDH1* | *NF1* | CDK2A |
|---|---|---|---|---|---|---|---|---|
| BTSC12 | M | 59Y | wt | mut | mut | wt | NA | NA |
| BTSC30 | M | 67Y | wt | wt | mut | wt | NA | NA |
| BTSC73 | M | 52Y | vIII | mut | mut | wt | NA | homo del |
| BTSC147 | M | 55Y | vIII | mut | mut | wt | wt | homo del |

**Supplementary Table. 3. Screening of common high confidence binding proteins for their impact on cell viability using an siRNA knockdown approach.** A pool of siRNA was designed to knockdown the genes encoding common high confidence binding proteins of OSMR. The sequences of sense siRNA and antisense siRNA for each target gene are provided.

| Target Gene Name | Sense siRNA Sequence | Antisense siRNA Sequence |
|---|---|---|
| B-cell receptor-associated protein 31 | UGGUGACUCUCAUUUCGCAtt | UGCGAAAUGAGAGUCACCAgg |
| B-cell receptor-associated protein 31 | GCAAGUUGGAUGUCGGGAAtt | UUCCCGACAUCCAACUUGCct |
| B-cell receptor-associated protein 31 | GCACUAAGCAAAAACUAGAtt | UCUAGUUUUUGCUUAGUGCtg |
| BCL2-like 14 (apoptosis facilitator) | CCGAGUAGCUGAAAUUGUUtt | AACAAUUUCAGCUACUCGGtt |
| BCL2-like 14 (apoptosis facilitator) | CCAUAGAAUUCAAAAUCCUtt | AGGAUUUUGAAUUCUAUGGtg |
| BCL2-like 14 (apoptosis facilitator) | AAGAGAUUUUUGUAACUGAtt | UCAGUUACAAAAAUCUCUUtg |
| small integral membrane protein 21 | GCAUAUUUCGAAACUUUCUtt | AGAAAGUUUCGAAAUAUGCtg |
| small integral membrane protein 21 | GCUGGGAACAUUUAAACAAtt | UUGUUUAAAUGUUCCCAGCtg |
| small integral membrane protein 21 | CACCAUAUUCGUUUCUUCAtt | UGAAGAAACGAAUAUGGUGtt |
| chromosome 8 open reading frame 44 | CAGAUGUACCGAGUUUGAAtt | UUCAAACUCGGUACAUCUGgt |
| chromosome 8 open reading frame 44 | CCGAGUUUGAAAUCCCAGAtt | UCUGGGAUUUCAAACUCGGta |
| chromosome 8 open reading frame 44 | GUAUUAGAAUUAUUUCUGAtt | UCAGAAAUAAUUCUAAUACct |
| C-type lectin domain family 2, member B | GAAUUUUCUUAGGCGGUAUtt | AUACCGCCUAAGAAAAUUCat |
| C-type lectin domain family 2, member B | AUACAACUGUUCCACUCAAtt | UUGAGUGGAACAGUUGUAUtt |
| C-type lectin domain family 2, member B | CAGCAACAGCUAGAUGUUAtt | UAACAUCUAGCUGUUGCUGca |
| chloride intracellular channel 1 | GGUUUUAGACAAUUACUUAtt | UAAGUAAUUGUCUAAAACCtt |
| chloride intracellular channel 1 | CAGCUGGGCUGGACAUAUUtt | AAUAUGUCCAGCCCAGCUGtg |
| chloride intracellular channel 1 | GUUUUAGACAAUUACUUAAtt | UUAAGUAAUUGUCUAAAACct |
| cAMP responsive element binding protein 3 | CUGUCUCUAUGGAUCUAGAtt | UCUAGAUCCAUAGAGACAGtt |
| cAMP responsive element binding protein 3 | GGCUAGUACUGACAGAUGAtt | UCAUCUGUCAGUACUAGCCta |
| cAMP responsive element binding protein 3 | GAGUGAGAGCUGUAGAAAAtt | UUUUCUACAGCUCUCACUCtc |
| chondroitin sulfate N-acetylgalactosaminyltransferase 2 | GGUCAUUAAUAAUCCUGAUtt | AUCAGGAUUAUUAAUGACCtc |
| chondroitin sulfate N-acetylgalactosaminyltransferase 2 | CUAGUGAUCUUUUAGAGUUtt | AACUCUAAAAGAUCACUAGgt |
| chondroitin sulfate N-acetylgalactosaminyltransferase 2 | GGACCUCUCAUGAAAGUGAtt | UCACUUUCAUGAGAGGUCCaa |
| SLIT and NTRK-like family, member 2 | CCAGUAGCCUAUUACCGAAtt | UUCGGUAAUAGGCUACUGGgt |
| SLIT and NTRK-like family, member 2 | GGAACCGUCUUGUCAAUGAtt | UCAUUGACAAGACGGUUCCat |
| SLIT and NTRK-like family, member 2 | GACUGUAUCCAAACGAAUUtt | AAUUCGUUUGGAUACAGUCtt |
| decorin | GCUGGACCGUUUCAACAGAtt | UCUGUUGAAACGGUCCAGCcc |
| decorin | AGAGGCUUAUUUGACUUUAtt | UAAAGUCAAAUAAGCCUCUct |
| decorin | GCUAGAUACUGGAAACCUAtt | UAGGUUUCCAGUAUCUAGCtt |
| eukaryotic translation initiation factor 2C, 3 | GAAGAGACAUCACACUCGAtt | UCGAGUGUGAUGUCUCUUCtg |
| eukaryotic translation initiation factor 2C, 3 | CAGUCGUCCUUCACACUAUtt | AUAGUGUGAAGGACGACUGgt |
| eukaryotic translation initiation factor 2C, 3 | CCAUAUGAGUUCGAUUUUUtt | AAAAAUCGAACUCAUAUGGgt |

| | | |
|---|---|---|
| group-specific component (vitamin D binding protein) | GCUUAAACAUUUAUCACUUtt | AAGUGAUAAAUGUUUAAGCtg |
| group-specific component (vitamin D binding protein) | CUACCUGUUUUAAUGCUAAtt | UUAGCAUUAAAACAGGUAGtt |
| group-specific component (vitamin D binding protein) | GAUCCAAAGGAAUAUGCUAtt | UAGCAUAUUCCUUUGGAUCtt |
| glutaredoxin (thioltransferase) | CAACCACACUAACGAGAUUtt | AAUCUCGUUAGUGUGGUUGgt |
| glutaredoxin (thioltransferase) | GCAGUGAUCUAGUCUCUUUtt | AAAGAGACUAGAUCACUGCat |
| glutaredoxin (thioltransferase) | GAGUCUUUAUUGGUAAAGAtt | UCUUUACCAAUAAAGACUCga |
| GC-rich promoter binding protein 1 | GCAUGGACAGAGAAUCGUUtt | AACGAUUCUCUGUCCAUGCaa |
| GC-rich promoter binding protein 1 | GGAAGUUCCCGUUCUCGUAtt | UACGAGAACGGGAACUUCCac |
| GC-rich promoter binding protein 1 | GAAAGGGAUAUAAACCGAAtt | UUCGGUUUAUAUCCCUUUCtt |
| histone cluster 1, H3a | CUGAACUGCUUAUUCGUAAtt | UUACGAAUAAGCAGUUCAGtg |
| histone cluster 1, H3a | GUAGGGCUAUUUGAGGACAtt | UGUCCUCAAAUAGCCCUACca |
| histone cluster 1, H3a | GCAAACAGUUGGCCACUAAtt | UUAGUGGCCAACUGUUUGCgt |
| IMP2 inner mitochondrial membrane peptidase-like (S. cerevisiae) | GUGGUGACAUUGUAUCAUUtt | AAUGAUACAAUGUCACCACgg |
| IMP2 inner mitochondrial membrane peptidase-like (S. cerevisiae) | GGGUUGAAGGUGAUCAUCAtt | UGAUGAUCACCUUCAACCCag |
| IMP2 inner mitochondrial membrane peptidase-like (S. cerevisiae) | AGAGAGUGAUUGCUCUUGAtt | UCAAGAGCAAUCACUCUCUta |
| leucine-rich repeats and IQ motif containing 1 | GGUCUUUGUGAUACACCUAtt | UAGGUGUAUCACAAAGACCtt |
| leucine-rich repeats and IQ motif containing 1 | CAGCUUGACUAAAAUCGUAtt | UACGAUUUUAGUCAAGCUGtt |
| leucine-rich repeats and IQ motif containing 1 | CUACCAUUGUAUACCUAGAtt | UCUAGGUAUACAAUGGUAGgt |
| N-acetyltransferase 8 (GCN5-related, putative) | CAGUCUUUUUGAUUCCCAUtt | AUGGGAAUCAAAAAGACUGaa |
| N-acetyltransferase 8 (GCN5-related, putative) | GCUUAUUGUCCAUUCACAUtt | AUGUGAAUGGACAAUAAGCtt |
| N-acetyltransferase 8 (GCN5-related, putative) | CAACCUUUCACUGCAAUGAtt | UCAUUGCAGUGAAAGGUUGgg |
| nurim (nuclear envelope membrane protein) | CCUUCUCGUCUUUGACUAUtt | AUAGUCAAAGACGAGAAGGat |
| nurim (nuclear envelope membrane protein) | CGGGCCCAGCUACAAAGAAtt | UUCUUUGUAGCUGGGCCCGga |
| nurim (nuclear envelope membrane protein) | GCCUCAAACAGGUAUACUAtt | UAGUAUACCUGUUUGAGGCcc |
| phosphatidylinositol glycan anchor biosynthesis, class K | GGACAUCGCACUGAUCUUUtt | AAAGAUCAGUGCGAUGUCCag |
| phosphatidylinositol glycan anchor biosynthesis, class K | GGCUCUAGCUAGUAGUCAAtt | UUGACUACUAGCUAGAGCCat |
| phosphatidylinositol glycan anchor biosynthesis, class K | CCAACAUAGAACUCGCGGAtt | UCCGCGAGUUCUAUGUUGGta |
| psoriasis susceptibility 1 candidate 1 | AGAAGUAACAUGUCCAAAAtt | UUUUGGACAUGUUACUUCUga |
| psoriasis susceptibility 1 candidate 1 | UCCAAGCAAUGAUAUCCAAtt | UUGGAUAUCAUUGCUUGGAgg |
| psoriasis susceptibility 1 candidate 1 | GCAAUGAUAUCCAAGGAAUtt | AUUCCUUGGAUAUCAUUGCtt |
| phosphatidylserine synthase 1 | GGAUGAUGUGAACUACAAAtt | UUUGUAGUUCACAUCAUCCtt |
| phosphatidylserine synthase 1 | GAGUGUACCUUUUCAUGAUtt | AUCAUGAAAAGGUACACUCca |
| phosphatidylserine synthase 1 | CCAUCGUCAGCCUCAUGUAtt | UACAUGAGGCUGACGAUGGtg |
| RAB1B, member RAS oncogene family | GCUGAAAUCAAAAAGCGGAtt | UCCGCUUUUUGAUUUCAGCag |
| RAB1B, member RAS oncogene family | GCACCAGCCUUAACCCUCAtt | UGAGGGUUAAGGCUGGUGCcc |
| RAB1B, member RAS oncogene family | AGAGCGACCUCACCACCAAtt | UUGGUGGUGAGGUCGCUCUtg |

| | | |
|---|---|---|
| SET domain containing (lysine methyltransferase) 7 | GGACCGCACUUUAUGGGAAtt | UUCCCAUAAAGUGCGGUCCtc |
| SET domain containing (lysine methyltransferase) 7 | GCACGUAUGUAGACGGAGAtt | UCUCCGUCUACAUACGUGCcc |
| SET domain containing (lysine methyltransferase) 7 | CCUGGACGAUGACGGAUUAtt | UAAUCCGUCAUCGUCCAGGtg |
| solute carrier family 35, member B3 | CCUACAUGAUAAUAGCUUUtt | AAAGCUAUUAUCAUGUAGGtt |
| solute carrier family 35, member B3 | CUGUAUGAUUUGAUAAACAtt | UGUUUAUCAAAUCAUACAGtg |
| solute carrier family 35, member B3 | GGACCUAUGGUUAUGCGUUtt | AACGCAUAACCAUAGGUCCga |
| ST3 beta-galactoside alpha-2,3-sialyltransferase 4 | GCAGACCAUUCACUACUAUtt | AUAGUAGUGAAUGGUCUGCtt |
| ST3 beta-galactoside alpha-2,3-sialyltransferase 4 | UCUGGGAUGUCAAUCCUAAtt | UUAGGAUUGACAUCCCAGAtg |
| ST3 beta-galactoside alpha-2,3-sialyltransferase 4 | GGAAGACAGUUUUUAUUUUtt | AAAAUAAAAACUGUCUUCCcg |
| TRAF family member-associated NFKB activator | CACUCAAGAUAACAAUUAUtt | AUAAUUGUUAUCUUGAGUGga |
| TRAF family member-associated NFKB activator | GAACUAUGAGCAGAGAAUAtt | UAUUCUCUGCUCAUAGUUCtc |
| TRAF family member-associated NFKB activator | GAGAUUCUGCAGUAAAAGAtt | UCUUUUACUGCAGAAUCUCta |
| transmembrane protease, serine 6 | GGAACUUACUACAACUCCAtt | UGGAGUUGUAGUAAGUUCCca |
| transmembrane protease, serine 6 | GCCUGUGAAGUGAACCUGAtt | UCAGGUUCACUUCACAGGCct |
| transmembrane protease, serine 6 | GCUGACCGCUGGGUGAUAAtt | UUAUCACCCAGCGGUCAGCga |
| zona pellucida glycoprotein 1 (sperm receptor) | CAGUCUUCUCGGCCGAUUAtt | UAAUCGGCCGAGAAGACUGca |
| zona pellucida glycoprotein 1 (sperm receptor) | ACACUGGGAUGUGAACAAAtt | UUUGUUCACAUCCCAGUGUtc |
| zona pellucida glycoprotein 1 (sperm receptor) | CACCCACUGUGGAACCACAtt | UGUGGUUCCACAGUGGGUGag |

**Supplementary Table. 4. Generation of CLIC1 CRIPSR transgenic BTSCs**. Two different gRNA was employed to generate CLIC1-CRISPR monoallelic and biallelic deletions in BTSC73 and BTSC147. The sequences for the gRNAs as well as the primers employed to screen each clone are listed.

| | |
|---|---|
| gRNA1-CLIC1-Fwd | caccGTCAACGGTGGTAACATTGA |
| gRNA1-CLIC1-RC | aaacTCAATGTTACCACCGTTGAC |
| gRNA2-CLIC1-Fwd | caccGTACCGATGCACTCCCCGGA |
| gRNA2-CLIC1-RC | aaacTCCGGGGAGTGCATCGGTAC |
| CLIC1 Internal-Fwd | TAGCTGAGGTTCCTCCCAGG |
| CLIC1 Internal-Rev | TATTCCTCCCAGGACCCAGG |
| CLIC1 External-Fwd | AGGGACTGGCCTAGGGATG |
| CLIC1 External-Rev | AAAATGGAGGGGGTTGAGGG |

**Supplementary Table. 5.** Primer sequences for OSMR, CLIC1 and the housekeeping genes, GUSB and ACTB, are provided.

| Gene | Forward Primer Sequence | Reverse Primer sequence |
|---|---|---|
| GUSB | GCGTTCCTTTTGCGAGGAGA | GGTGGTATCAGTCTTGCTCAA |
| ACTB | CAGCAGATGTGGATCAGCAAG | GCATTTGCGGTGGACGAT |
| OSMR | ACTGGAACCTGCCACAGAGT | TCCAAGCTCACAATTCTCCA |
| CLIC1 | ACCGCAGGTCGAATTCTTC | ACGGTGGTAACATTGAAGGTG |

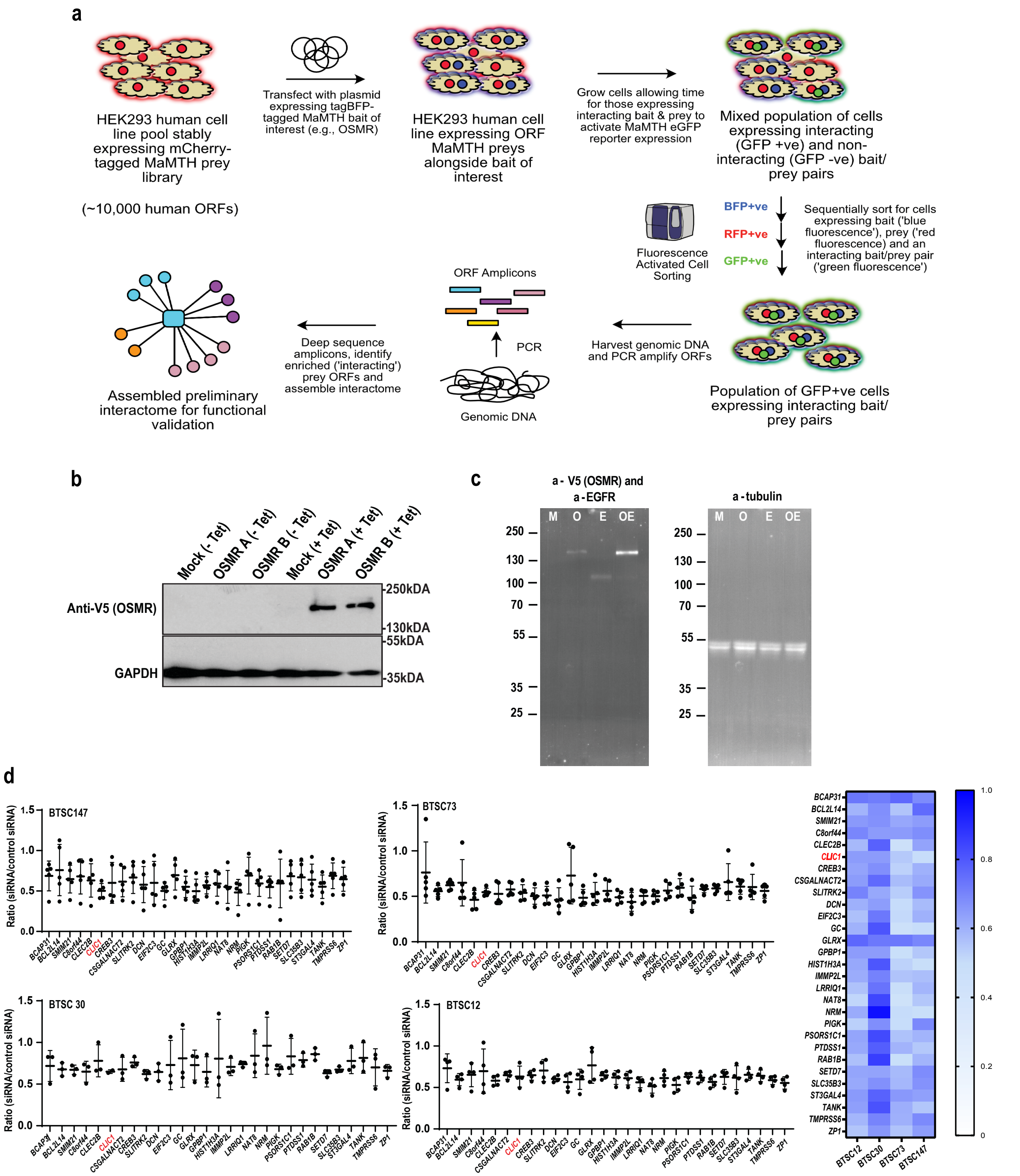


**Supplemental Fig. 1. Schematic outline of the MaMTH-HTS screening workflow and mapping global binding partners of OSMR and Characterization of OSMR bait induction in MaMTH HEK293T reporter cells.**

**a**, Expressed bait and prey are indicated by blue circles (corresponding to tagBFP tag) and red circles (corresponding to mCherry tag), respectively. Green circles (corresponding to expressed

GFP reporter) are indicative of bait/prey interaction. **b**, Western blot analysis of HEK293T reporter cells transfected with OSMR MaMTH bait construct verifying that tagged OSMR is properly expressed in response to induction with 0.5 mg/ml tetracycline, with no detectable leakage in the absence of inducer. 'OSMR A' and 'OSMR B' correspond to MaMTH bait constructs tagged at their C-terminus with Cub-GAL4TF-V5 or Cub-GAL4TF-V5-P2A-tagBFP, respectively. GAPDH was used as a loading control. **c,** OSMR and EGFRvIII proof of expression in the bait. **d**, Cell viability was assessed in four different patient-derived BTSCs (#12, #30, #73, and #147), transfected with siRNAs targeting different genes encoding candidate binding proteins. Each graph is normalized to the control group transfected with a non-targeting siRNA. Data are presented as means ± SEM, n=3 biological replicates in a-d with Heatmap plot comparing the differences across four BTSCs. The colour gradient, represented by column Z-scores, illustrates relative cell viability assessed by PrestoBlue analysis.

**M**: Mock transfected, **O**: OSMR Bait transfected, **E**: EGFRvIII transfected, **OE**: OSMR Bait and EFGRvIII co-transfected.

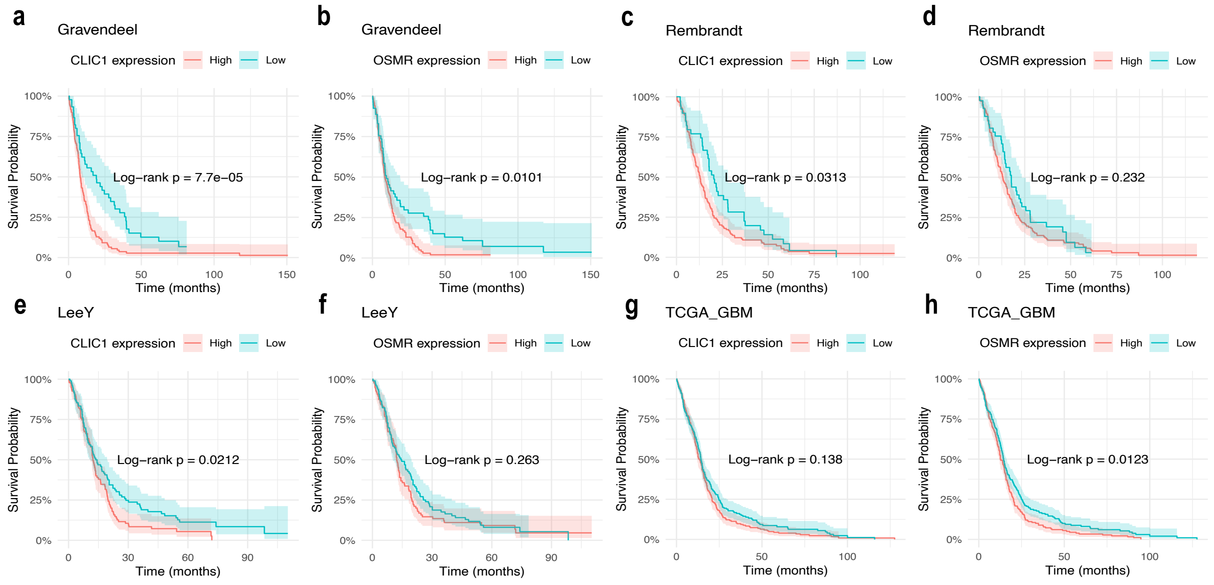


**Supplemental Fig. 2. Clinical relevance of expression of CLIC1 and OSMR using public datasets on patients' survival**.

Kaplan–Meier survival curves for glioblastoma patients stratified by median gene expression of CLIC1 and OSMR in Gravendeel (**a**,**b**), Rambrandt (**c**,**d**), LeeY (**e**,**f**), and TCGA_GBM (**g**,**h**) datasets. Rose lines represent patients with high gene expression (> median), while turquoise lines indicate low expression (< median). Across Gravendeel, Rambrandt, and LeeY datasets, higher CLIC1 expression is associated with reduced survival probability over time, whereas the impact of OSMR expression on survival is less consistent.

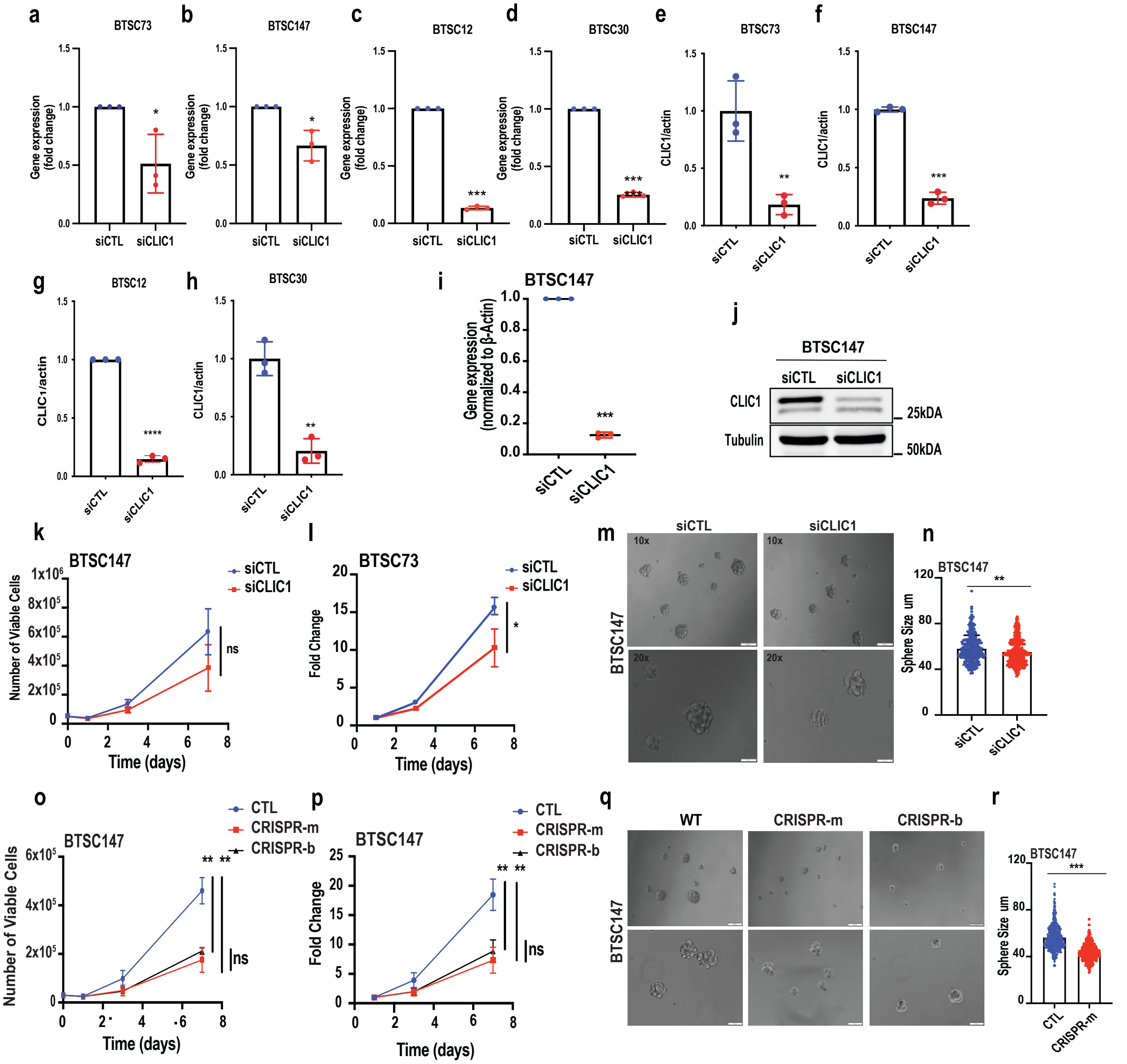


**Supplemental Fig. 3**. **Effects of CLIC1 gene modification on BTSC viability and sephere size. a-d**, BTSC73, BTSC147, BTSC12, and BTSC30 cells were transfected with an siRNA pool targeting CLIC1 (siCLIC1) or non targeting siRNA control (siCTL) and mRNA expression of CLIC1 in each BTSC line was assessed by RT-qPCR. Data was normalized to the housekeeping gene, GUSB. **e-h**, BTSCs were subjected to immunoblotting analysis using an antibody to CLIC1 and the CLIC1 protein expression level was normalized to ACTIN, n=3 biological replicates. Statistical analysis was performed using a student t-test. $**P < 0.01$, $***P < 0.001$, $***P < 0.0001$. **i**, BTSC147 was transfected with an siRNA targeting CLIC1 (siCLIC1) or non targeting siRNA control (siCTL) and mRNA expression of CLIC1 was assessed by RT-qPCR. Data was normalized to the housekeeping gene, GUSB. **j**, The protein expression of CLIC1 was assessed by immunoblotting using a CLIC1antibody. Tubulin was used as a loading control. **k,l**, Proliferation ability plotted based on the number of viable cells counted using trypan blue exclusion dye (**k**). ell viability was plotted as fold change based on the initial count on day1 for each condition (**l**). A student-t test was used to compare two conditions at day 7. Data are presented as mean ± SD, n = 3 biological replicates, non-significant (ns), *$P < 0.05$. **m**, Representative images of siCLIC1 and siCTL sphere sizes following 3 days of incubation at 37°C and 5% CO2. Sphere images were taken using 10x (top panel, scale bar 100 µm) and 20x (bottom panel, scale bar 100 µm) objectives. **n**, Sphere size was quantified. A minimum of 100 spheres was assessed from each biological replicate and data was plotted as a mean ± SD. A student-t test was used to identify significance, **$P < 0.01$. **o-r**, Two different transgenic CLIC1 CRISPR lines were generated for each of BTSC147 and BTSC73. **o**, Proliferation ability plotted based on the number of viable cells counted using trypan blue exclusion dye. **p**, Proliferation capability plotted as fold change based on the initial count on day1 for each condition. A student-t test was used to compare two conditions at day 7. Data are presented as mean ± SD, n=3 biological replicates, non-significant (ns), **$p<0.01$. **q**, Representative images of CLIC1-CRISPR cells and control BTSC147 sphere sizes following 3 days of incubation at 37°C and 5% CO2. Sphere images were taken using 10x (top panel, scale bar 100 µm) and 20x (bottom panel, scale bar 100 µm) objectives. **r**, Sphere sizes in CRISPR and control BTSC147 were quantified. A minimum of 100 spheres was assessed from each biological replicate and data was plotted as a mean ± SD. A student-t test was used to identify significance, ***$P < 0.001$.

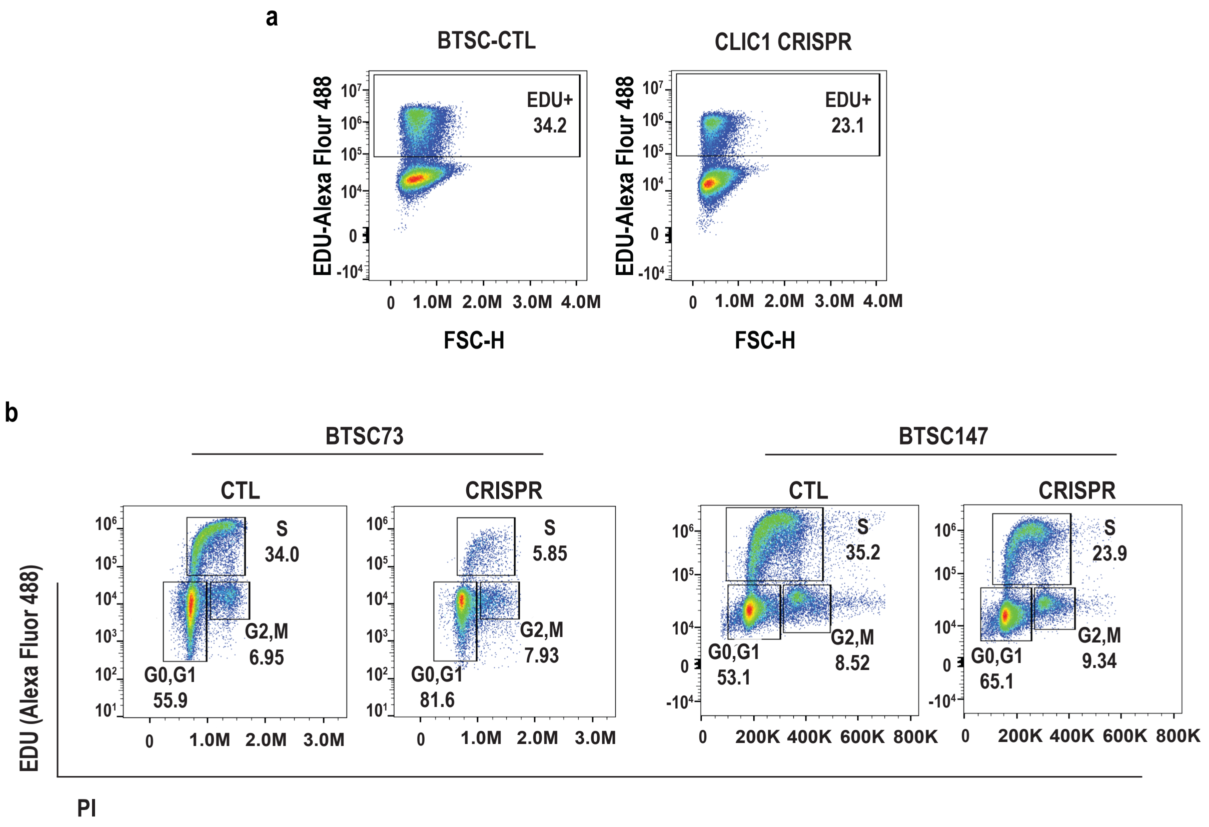


**Supplemental Fig. 4**. **Cell cycle analysis in EdU-Labeled Cells.**

**a**, BTSC147 cells and their related CLIC1 CRISPR line were labeled with the Click-iT EdU-Alexa 488 kit, and the samples were then analyzed using a flow cytometer. **b**, BTSC73 and BTSC147 cells, along with their respective CLIC1 CRISPR lines, were simultaneously stained with EdU and propidium iodide (PI). The resulting dual-parameter plots, which combine DNA content and EdU incorporation data, show three distinct cell populations. This analysis allows for the direct quantification of cells in the S-phase, as well as the identification of the G0/G1 and G2/M populations.

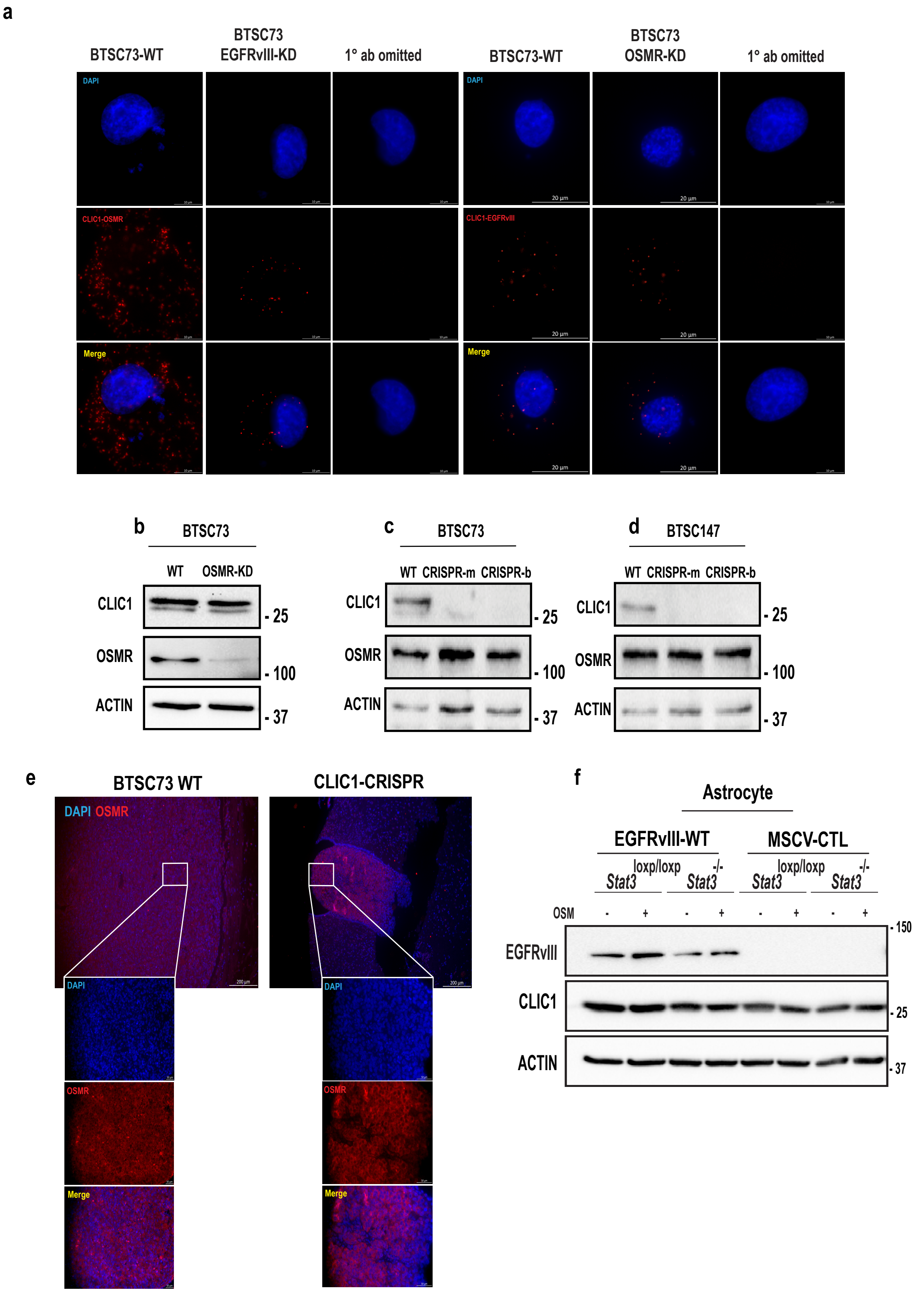


**Supplemental Fig. 5. Proximity ligation assay and Correlative protein analysis following OSMR and CLIC1 gene expression alternation.**

**a,** BTSC73-CTL, BTSC73 EGFRvIII-KD, and BTSC73 OSMR-KD were subjected to PLA analysis using antibodies to CLIC1 and OSMR, and CLIC1 and EGFRvIII. Each red puncta represents PLA signal. Primary antibodies were omitted for control group. Nuclei were stained with DAPI. **b**, BTSC73 cells were subjected to OSMR KD using siRNA, and the protein expression of CLIC1 was assessed by immunoblotting with an anti-CLIC1 antibody. Actin was used as a loading control. **c**, BTSC73 and **d**, BTSC147 cells, along with their related CLIC1 CRISPR lines, were subjected to Western blotting using an anti-CLIC1 antibody and an anti-OSMR antibody. Actin was used as a loading control. **e**, Brain tissues from the BTSC73 WT and CLIC1-CRISPR groups were subjected to immunostaining with an anti-OSMR antibody to detect the protein's localization and expression. Nuclei were counterstained with DAPI. **f**, EGFRvIII expressing *STAT3*$^{loxp/loxp}$, *STAT3*$^{-/-}$ and murine stem cell virus (MSCV)-CTL (non-EGFRvIII expressing) *STAT3*$^{loxp/loxp}$, *STAT3*$^{-/-}$ astrocytes[1], were treated with 10 ng/mL OSM for 5 minutes and subjected to immunoblotting using antibodies against EGFRvIII and CLIC1. ACTIN was used as a loading control.

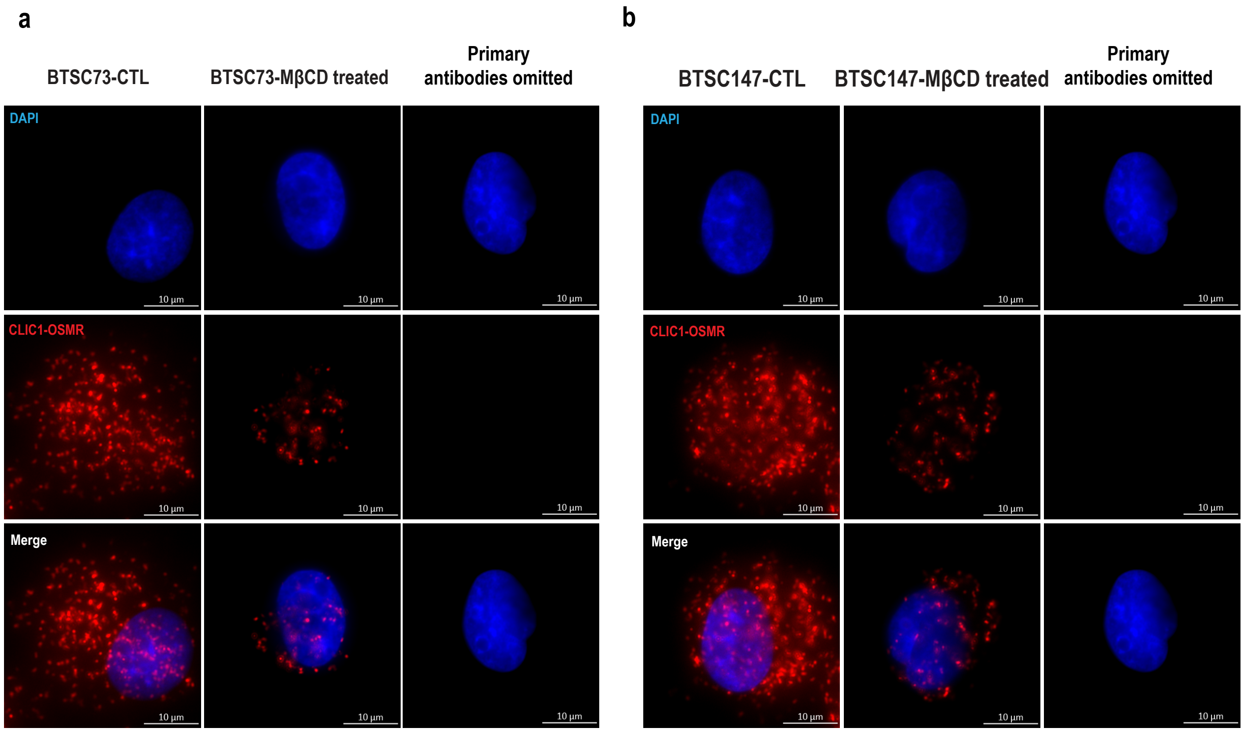


**Supplemental Fig. 6. Lipid Raft Disruption Reduces the Proximity of OSMR-CLIC1 in BTSCs.**

**a**, BTSC73 and **b**, BTSC147 cells were subjected to lipid raft disruption using 5 mM Methyl-β-cyclodextrin (MβCD) for 1 hour at 37°C. Subsequently, a proximity ligation assay (PLA) was performed using anti-CLIC1 and anti-OSMR antibodies. Each red puncta indicates a PLA signal. Primary antibodies were omitted for the control group. Cell nuclei were stained with DAPI. Scale bar, 10 μm.

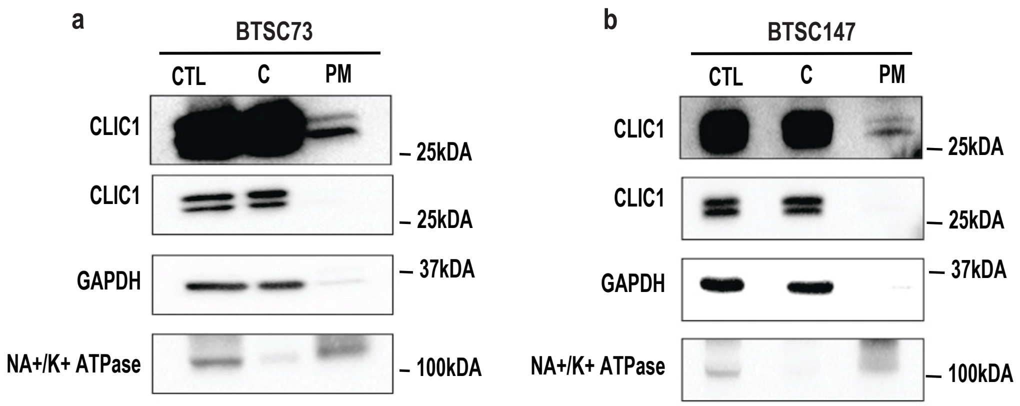


**Supplemental Fig. 7. Subcellular localization of CLIC1 in BTSC**.

**a**, BTSC73 and **b**, BTSC147 were subjected to subcellular fractionation and cytoplasmic, C, and plasma membrane, PM, fractions were analyzed by immunoblotting using the indicated antibodies. Whole cell lysates were loaded on the same gel. $Na^+/K^+$ ATPase and GAPDH were used as PM and cytoplasmic markers, respectively. High and low exposure images are shown for CLIC1 expression. n = 3 biological replicates.

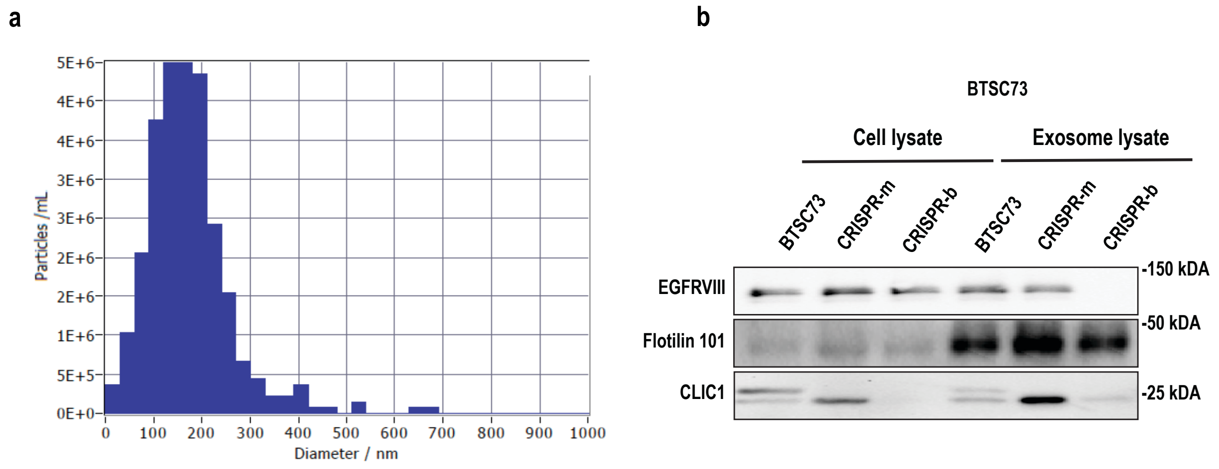


**Supplemental Fig. 8. Extracellular vesicles (EVs) isolation.**

BTSC73 cell supernatant was centrifuged at 200 x g for 10 minutes, 300 x g for 10 minutes, 2,000 x g for 10 minutes, 10,000 x g for 30 minutes, and 100,000 x g for 2 hours, respectively. After removing the supernatant and resuspending the EV pellet in PBS, the resulting pellet was then washed and centrifuged at 49,000 rpm for 30 minutes. **a**, Particle size distributions (PSDs) for BTSC73 were analyzed in the range of 0-1000 nm using ZetaView. **b**, EV and whole cell lysates fractions from CLIC1 CRISPRs and control BTSC73 were analyzed by immunoblotting using annotated antibodies.

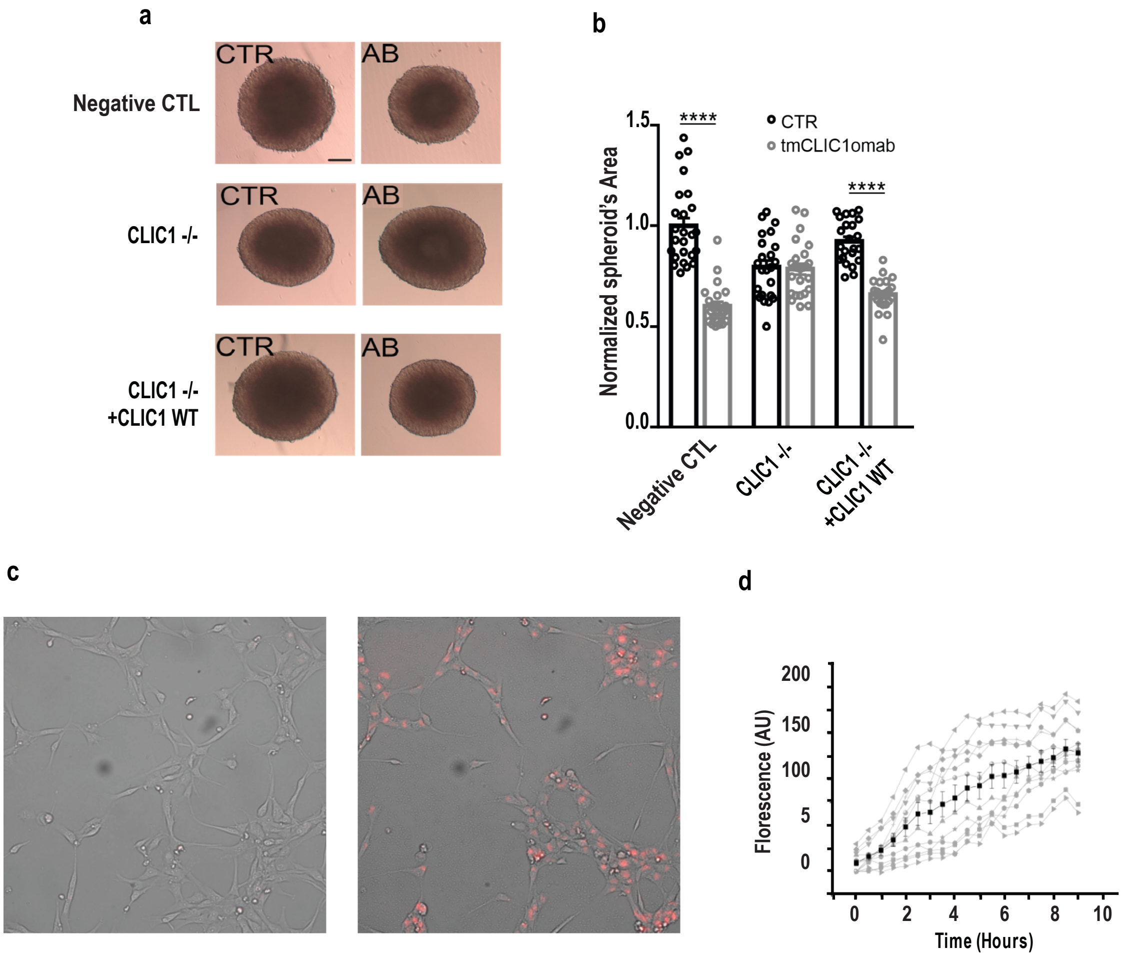


**Supplemental Fig. 9. tmCLIC1omab validation.**

Human glioblastoma primary cell line 3D culture in presence of tmCLIC1omab is shown. **a**, Representative images showing the development of GBM1 3D structure (spheroid) after 72h of incubation in the absence or presence of tmCLIC1omab as indicated in each square. **b**, Quantification of spheroid area in the absence (black circles) or presence (grey circles) of tmCLIC1omab. NC: CT, n=25, tmCLIC1omab n=23; ****p<0.0001. $Clic1^{-/-}$: CT n=25, tmCLIC1omab n=23. Clic1-/- + Clic1 WT: CT, n=23, tmCLIC1omab n=23; ****p<0.0001. Mean ± SEM, one-way ANOVA, Tukey's multiple comparison test. Scale bar 100 µm. **c**, Representative picture of control cells or cells incubated with tmCLIC1omab labeled with pHRodo red dye. **d**, Quantification of fluorescence deriving from labeled particles with tmCLIC1omab and the dye. The average level of fluorescence is represented by black line. Grey points represent the single well.

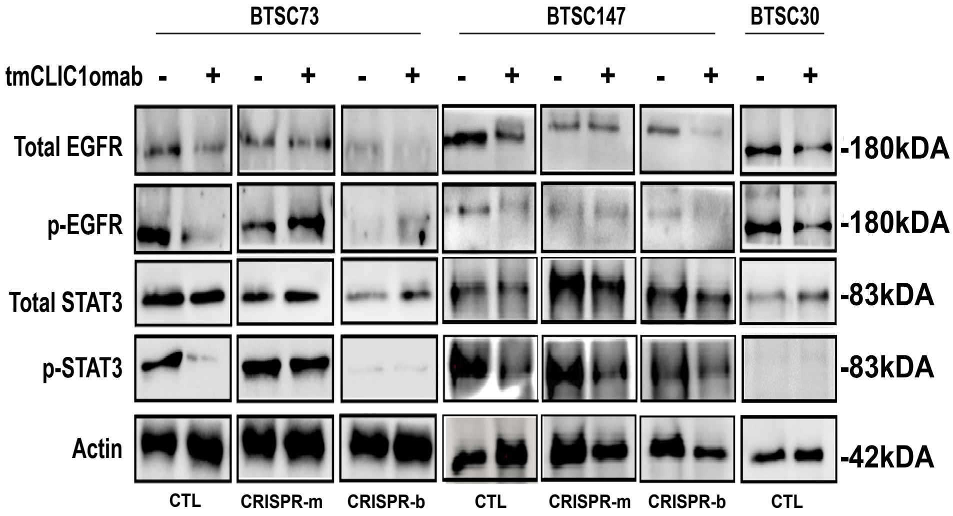


**Supplemental Fig. 10. Pharmacological inhibition of PM-CLIC1 attenuates the phosphorylation of STAT3 and EGFR.**

BTSC73, BTSC147, and BTSC30, as well as corresponding CLIC1-CRISPR BTSCs were treated with a monoclonal tmCLIC1mab or IgG control. The whole cell lysates were subjected to immunoblotting analysis using antibodies indicated on the blots. Actin was used as a loading control.

**Supplementary Material and methods**

**1. BTSC culture**

Prior to use in experiments, BTSC lines were thawed from a cryopreserved state (10% dimethyl sulfoxide -DMSO) and placed in BTSC media containing serum-free NeuroCult NS-A medium (StemCell Technologies, Inc., #05750), supplemented with 100 U/ml penicillin, 100 μg/ml streptomycin (Sigma Aldrich, P4333), heparin (2 μg/ml, StemCell Technologies, Inc., #07980), human EGF (20 ng/ml, Miltenyi Biotec, #130-093-825), and human FGF (10 ng/ml, Miltenyi Biotec, #130-093-838). Cultures were kept in T-75 or T-25 culture flask for cells in suspension (Sarstedt #833911502, 833910502) and grown as spheres in suspension. All cell lines were subject to analysis for the presence of mycoplasma using the PCR method. Passaging of BTSC spheres was done every 3-7 days or once spheres reached 200-250 μm in size. This was accomplished by collecting media containing BTSC spheres followed by centrifugation at 1000 x g for 10 mins. Cells were dissociated into a single cell suspension by incubating with 0.2 ml Accumax dissociation solution (Innovative Cell Technologies, #AM105) for 10 minutes at 37°C followed by the addition of 0.8 ml BTSC media. Utilizing Trypan Blue Stain 0.4% (Gibco, #15250-061), cells were counted and subsequently transferred onto low attachment flasks for culturing in BTSC suspension media. Cell plating densities were determined based on the size of the plating vessel (T-75 or T-25 flask) and the duration of the desired incubation (3-7 days).

**2. Cell population growth assay**

BTSC spheres were processed into a single-cell suspension using Accumax dissociation solution (Innovative Cell Technologies, #AM105). Single cells were plated at a density of 30,000 or 50,000 cells per well (6-well plate) in 2 ml BTSC media for CRISPR/CAS9 clones and siRNA experiments, respectively. 1 day (24 hours), 3 days (72 hours), and 7 days (168 hours) following plating, cells were collected, and live cells were assessed using trypan blue exclusion dye (Gibco #15250-061). This dye differentiates live and dead cells by the uptake or lack of uptake of a blue dye. Counts were done using an automated cell counter (Countess II FL Automated Cell Counter).

**3. Sphere size assessment**

Sphere size assessment was used to determine any differences in size between BTSC WT spheres, or spheres generated from either siRNA transient knockdown or CRISPR-CAS9 stable knockdown, or knockout. BTSC spheres were collected and processed into a single cell suspension using Accumax dissociation solution (Innovative Cell Technologies, #AM105). Cells were plated as follows: 3 to 6 wells containing 500, 250, and 125 cells respectively in 100 ul BTSC media in a 96-well plate. The purpose of doing this was to ensure that the plating density did not impact the size of the spheres. By maintaining consistent plating densities, potential confounding factors affecting sphere size could be mitigated, allowing for a more accurate and reliable assessment of experimental results. 72 hours following plating and incubation, sphere size was assessed, and visualizations captured using a scanning microscope (Olympus LS, IXplore Pro Automated Microscope system) equipped with the adjoining imaging software (Olympus LS, cellSens Version 1.12) using the 'Measure' feature at 10-20x objective magnification.

**4. Cell Fractionation**

The plasma membrane and cytoplasm were separated using the Thermofisher Subcellular Protein Fractionation Kit (Thermofisher, #78840), following the manufacturer's protocol. Briefly, $5 \times 10^7$ BTSCs were rinsed once with 1 ml of 1X PBS containing 0.3% BSA and then with 1 ml of 0.9% sodium chloride solution. For cytoplasm extraction, cells were treated with 500 µl of cytoplasmic extraction buffer (CEB) and shaken for approximately five minutes at 4°C. After centrifugation at 500 x g for 5 minutes, the supernatant was carefully collected. For plasma membrane extraction, the pellet was incubated with 500µl of membrane extraction buffer and shaken for approximately five minutes at 4°C. Subsequently, the cells were centrifuged at 3000 x g for 5 minutes, and the supernatant was collected.[2] Protein concentration of cytosolic, and plasma membrane lysates was determined using the Bradford assay (Bio-Rad, #5000006) and 10 ug of protein was loaded in a 10% SDS PAGE gel.

**5. Electroporation of BTSC**

BTSC electroporation was conducted in accordance with our established protocols.[3] Briefly, at a concentration of 100nM, siRNA was delivered to patient-derived BTSCs using electroporation as described by the AMAXA nucleofector 2b device, set at 1300 volts (Lonza, #AAB1001). Electroporated cells were plated in a T-75 culture flask in suspension containing BTSC media and incubated at 37°C and 5% $CO_2$. Plating for experiments took place 24 hours after electroporation while collection of cells for gene expression analysis by RT-qPCR and protein expression by Immunoblotting were done 72 hours after electroporation.

**6. Transcriptomic database analysis**

Six processed glioblastoma datasets, referred to as Ducray[4], Gravendeel[5], Kamoun[6], LeeY[7], Rembrandt[8], and TCGA_GBM[9], were downloaded from the GlioVis database[10], version

0.20. Datasets were analyzed in R version 4.4.1.[11] Boxplot P-values were calculated using Mann-Whitney U tests. Survival analysis log-rank P-values were calculated using the survival package version 3.7-0[12,13], in R.

**7. Gene expression analysis**

RNA was extracted from cells using the TRIzol method. In this method, pelleted cells are incubated with 1 ml TRIzol digestive reagent (Invitrogen, #15596026) at room temperature for 5 minutes. Next, 0.2 ml chloroform (Sigma Aldrich, #288306) was added to the samples and shaken vigorously for 20 seconds followed by a 2–3-minute incubation at room temperature (22-25°C). Samples were then centrifuged for 15 minutes at 12000 x g, 4°C after which 0.5 ml of the upper aqueous phase was added to a clean tube containing 0.5ml isopropanol. Samples were again incubated at room temperature (22-25°C) for 15 minutes followed by centrifugation at 12000 x g for 10 minutes, 4°C. The resultant supernatant was removed from each sample, and the RNA pellet was washed first with 75% Ethanol and 100% Ethanol. Each wash step was followed by centrifugation at 7500 x g for 5 mins (4°C). The supernatant was removed, and RNA samples were left to dry for 10 minutes at room temperature (22-25°C) in a sterile biological hood. The RNA pellet was dissolved in 20-50 µl RNase-free water and incubated at 55-60°C for 10-15 minutes to ensure RNA dissolution. cDNA was obtained by reverse transcription using the 5X All-In-One RT MasterMix cDNA synthesis system (abm, #G592). Gene expression data was obtained from RT-qPCR analysis using the fluorescent dye SYBR Green (Biorad, #1725271) and primers, as described in **Supp. Table 5**, by combining 1µl of 5 mM forward primer, 1 µl 5 mM of the reverse primer and 5 µl SYBR Green to make the PCR mix. This mix was used to generate a final plated mix of 3µl of 30ng cDNA and 7µl PCR mix that was run on the QuantStudioTM 7 Flex Real-Time PCR System (Applied Biosystems). mRNA expression levels were normalized to one of two housekeeping genes: beta-glucuronidase (*GUSB*) and beta-actin (*ACTB*).

**8. Lipid raft disruption**

Lipid raft disruption with Methyl-β-cyclodextrin (MβCD) began by preparing a 5 mM MβCD solution (Selleckchem # S6827) in serum-free cell culture media and warming it to 37°C. Next, the medium was removed, and the cells were washed gently with 1X PBS. The pre-warmed 5 mM MβCD solution was added to the cells, followed by incubation at 37°C for 1 hour to deplete cholesterol from the cell membrane and disrupt lipid rafts. After the incubation period, the MβCD solution was removed, and the cells were washed with 1X PBS to eliminate any remaining MβCD. Fresh, complete cell culture medium was then added to the cells.

**9. Immunoblotting and antibodies**

Protein expression was analyzed by SDS PAGE as previously described[3]. Total protein from samples was harvested from a minimum of $5x10^5$ using 1X RIPA Buffer (Thermo Fisher Scientific, #89900) supplemented with protease and phosphatase inhibitors (Thermo Fisher Scientific, #A32959). Following cell lysis, protein concentration was assessed using Bradford Assay (Bio-Rad), after which samples were run on a 10-15% SDS-PAGE gel and transferred on a Nitrocellulose Membrane 0.45um (Bio-Rad, #1620115) for 1 hour and 30 minutes at 100 volts. Membranes were blocked for non-specific binding using 5% bovine serum albumin (BSA) in 1X TBST for 1-2 hours. Subsequently, the membranes were incubated with primary antibodies at 4°C overnight, followed by horseradish peroxidase (HRP)-conjugated secondary antibodies in a 5% bovine serum albumin (BSA) blocking solution. Proteins of interest were visualised by ClarityTM Western ECL Substrate (BioRad, 170-5060) using ChemiDoc Imaging System (Biorad). The antibodies used for immunoblotting include anti-OSMR (1:100, Santa Cruz Biotechnology, sc-271695, mouse), anti-CLIC1 (1:200, Santa Cruz Biotechnology, sc-81873, mouse), α-Tubulin (1:5000, Abcam, ab4074, mouse), HRP-conjugated secondary antibody (1:5000, BioRad, 1706516, mouse).

**10. Extreme Limiting Dilution Assay (ELDA)**

A single cell suspension and followed by dilution using BTSC complete media to generate a final plated density of 25, 12, 6, 3 and 1 cell per well. 100μl of the diluted single-cell suspensions was plated in a 96-well plate as follows: 12 wells plated for dilution 25, 12, 6, and 3 cells per well and 48 wells (4 replicates of 12 wells) plated for 1 cell per well dilution.[14] This was done for 1 cell per well plating to achieve comparable results as many times single cell death can occur following plating. Plates were incubated at 37 °C and 5% $CO_2$ for 7 days after which sphere formation was assessed. Analysis of the responding wells as a function of the plated wells was undertaken and data was input into an online software available at https://bioinf.wehi.edu.au/software/elda/. Using this data provided by the software, the number of cells required to generate a positive well (having a sphere) was determined for each condition. Stem cell frequency (SCF) or the probability of finding a stem cell in the bulk sample of cells was also given as a value of 1/stem cell frequency. To determine percent SCF 1/stem cell frequency was multiplied by 100.

### 11. Duolink Proximity Ligation Assay (PLA)

Single cell BTSC suspension was plated on Nunc Lab-Tek, II CC2 Chamber Slide System (Thermo Scientific, #154941) that was coated overnight with a Poly-D-Lysine (PDL) coating. The working PDL solution (10 µg/ml) was made from a 2 mg/ml stock (Fisher Scientific, # CB-40210) using sterile water and incubated on the chamber slide overnight at 37 °C and washed thoroughly (3 times) with sterile water prior to addition of cell suspension. Following Accumax (Innovative Cell Technologies, #AM105) dissociation the single cell suspension was plated at a density of ~$2.5x10^4$ cells in media containing 10% FBS for 1 to 6 hours until cells had visible projections. Following incubation, cells were quickly washed with 1X PBS and fixed using 4% paraformaldehyde for 15 min at room temperature. Next, samples were washed with 1X PBS and permeabilized using 0.5% Triton X-100 for 25 mins and subsequently blocked for non-specifying binding for 1 hour using the Duolink Blocking Solution provided in the starter kit with agitation at 37 °C. Samples were then incubated overnight at 4 °C in a humid chamber with primary antibodies against the protein of interest: anti-OSMR (1:400, Abnova, H00009180-D01P, rabbit) and anti-CLIC1 (1:400, Santa Cruz Biotechnology, sc-81873, mouse). Following overnight incubation, samples were washed with Wash Buffer A (available in the Starter Kit) and incubated with the PLUS and MINUS oligonucleotide probes conjugated to secondary antibodies for 1 h at 37 °C in a humid chamber. Samples were subsequently washed with Wash Buffer A and incubated with Ligation Buffer with Ligase (available in the Starter Kit) for 30 minutes at 37 °C in a humid chamber. Rolling Circle Amplification (RCA) was achieved by washing samples with Wash Buffer A and incubating with polymerase and amplification solution (available in the Starter Kit) containing nucleotides for 100 mins at 37 °C in a humid chamber and protected from light. Following RCA, samples were washed with Wash Buffer B and mounted with Duolink *In Situ* Mounting Medium (DUO82040, Sigma). PLA signals were visualized and captured by a laser scanning microscope (Olympus LS, IXplore Pro Automated Microscope system) equipped with the adjoining imaging software (Olympus LS, cellSens Version 1.12) at 40-60X objective magnification.

### 12. Expanded Version of tmCLIC1 Monoclonal Antibody Design and Generation

#### 12.1. Growth Curve

Cells ($2x10^4$) were plated and counted over time using Trypan Blue exclusion and an automated counter in the presence or absence of 100 μM of IAA94 tmCLIC1 inhibitor used as control or 3 μg/ml of tmCLIC1omab antibody. Triplicates were used per condition (**Fig. 6l,m**).

**12.2. Dose-response**

Cells were seeded in 96-well plates at a density of $2x10^4$/well and allowed to adhere overnight. The following day, cells were treated with increasing concentrations of tmCLIC1omab ranging from 0.1 to 3 μM. After 72 hours of treatment, cell viability was assessed using the MTT assay (**Fig. 6n**).

**12.3. 3D culture**

Human glioblastoma cells Negative Control (NC), knockout for clic1 ($Clic1^{-/-}$), and rescued with CLIC1 WT were cultured in three-dimensional spheroids using the hanging drop method. To generate spheroids, 20–30 μL droplets containing (500–2,000 cells/drop) were pipetted onto the inner surface of a sterile Petri dish lid. The lid was then inverted and placed over a dish containing sterile phosphate-buffered saline (PBS) or distilled water to maintain humidity and prevent evaporation. Plates were incubated at 37°C with 5% $CO_2$ for 24 hours, allowing spheroids to form through gravity-driven cell aggregation. After formation, spheroids were harvested by gently flushing the drops into a sterile conical tube using a pipette and transferred into low-attachment plates covered by 1,5% (w/v) Agarose for further analysis or treatment. Spheroids were transferred in complete media and /or to tmCLIC1omab-containing media. Images were captured at 0 and 72 h, and the spheroid area was quantified using ImageJ, and the area obtained after 72 hours on incubation was normalized on the initial spheroid's area (**Supp. Fig.9a,b**).

**12.4. tmCLIC1omab irreversible binding**

These experiments were performed as a growth curve analysis of human primary glioblastoma cell lines in which tmCLIC1omab was added for 3, 5, and 24 hours respectively and then washout and substituted with fresh medium. Proliferation was evaluated after 72 hours (**Fig. 6o**).

**12.5. tmCLIC1omab internalization**

pHrodo™ (Life Technologies; Catalog number P35372) was prepared according to the manufacturer's instructions, labeling the dye with tmCLIC1omab. Labeled particles were resuspended in warm assay buffer then added to the cells. Cells were incubated at 37°C for 30 min to allow uptake. In control wells, parallel incubations were done in the presence of antibody isotype to assess non-specific or blocked uptake. After incubation, cells were washed gently with PBS to remove excess particles and fluorescence microscopy was performed using a Spinning disk confocal microscope (Ex 560 nm/Em 585 nm). Quantification of fluorescence intensity was performed using ImageJ/Fiji. Increased fluorescence intensity indicates successful internalization and acidification of pHrodo-labeled particles within endosomes or lysosomes, as pHrodo becomes highly fluorescent in acidic environments but remains non-fluorescent at neutral pH (**Supp. Fig.9c,d**).

#### 12.6. Antibody sequences of tmCLIC1omab

##### 12.6.1. Heavy chain: DNA sequence (411 bp)

Signal sequence-FR1-CDR1-FR2-CDR2-FR3-CDR3-FR4

ATGAACTTCGGGCTCAGCTTGATTTTCCTTGTCCTTGTTTTAAAAGGTGTCCAGTGTG
AAGTGATGCTGGTGGAGTCTGGGGGAGGCTTAGTGAAGCCTGGAGGGTCCCTGAAA
CTCTCCTGTGCAGCCTCTGGATTTGCTTTCAGTAATTATGCCATGTCCTGGATTCGCC
AGATTCCGGAGAAGAGGCTGGAGTGGGTCGCAACCATTAGTACTGGTGGTAGTTCC
ACCTTCTATCCAGACAGTGTGAAGGGGCGATTCACCATCTCCAGAGACAATGCCAA
GAATACCCTATACCTGCAAATGAGCAGTCTGAGGTCTGAGGACACGGCCATGTATTA
CTGTACAAGACATGCCTACGAGGCCTGGTTTGCTTACTGGGGCCAAGGGACTCTGGT
CACTGTCTCTGCA

##### 12.6.2. Heavy chain: Amino acid sequence (137 aa)

Signal peptide-FR1-CDR1-FR2-CDR2-FR3-CDR3-FR4

MNFGLSLIFLVLVLKGVQCEVMLVESGGGLVKPGGSLKLSCAASGFAFSNYAMSWIRQI
PEKRLEWVATISTGGSSTFYPDSVKGRFTISRDNAKNTLYLQMSSLRSEDTAMYYCTRH
AYEAWFAYWGQGTLVTVSA

##### 12.6.3. Light chain: DNA sequence (384 bp)

Signal sequence-FR1-CDR1-FR2-CDR2-FR3-CDR3-FR4

ATGGATTTTCAAGTGCAGATTTTCAGCTTCCTGCTAATGAGTGCCTCAGTCATAATGT
CCAGGGGACAAATTGTTCTCACCCAGTCTCCAGCACTCATGTCTGCATCTCCAGGGG
AGAAGGTCACCATGACCTGCAGTGCCAGCTCAAGTGTAAGTTACATGTACTGGTACC
AACAGAGGCCAAGATCCTCCCCCAAACCCTGGATTTATCTCACATCCACCCTGGCTT
CTGGAGTCCCTACTCGCTTCAGTGGCAGTGGGTCTGGGACCTCTTACTCTCTCACAAT

CAGCAGCATGGAGGCTGAAGATGCTGCCACTTATTACTGCCAGCAGTGGAATAGTC
ACCCACTCACGTTCGGCTCGGGGACAAAGTTGGAAATAAAA

#### 12.6.4. Light chain: Amino acid sequence (128 aa)

Signal peptide-FR1-CDR1-FR2-CDR2-FR3-CDR3-FR4

MDFQVQIFSFLLMSASVIMSRGQIVLTQSPALMSASPGEKVTMTCSASSSVSYMYWYQQ
RPRSSPKPWIYLTSTLASGVPTRFSGSGSGTSYSLTISSMEAEDAATYYCQQWNSHPLTF
GSGTKLEIK

#### 12.7. Project Specifications

| Sequence | Clones sequenced | Clones with >99% sequence identity |
|---|---|---|
| $V_H$ | 5 | 5 |
| $V_L$ | 5 | 5 |

#### 12.8. international ImMunoGeneTics information system (IMGT) Analysis of V(D)J Junctions

| Sequence | V-GENE and allele | Functionality | V-REGION identity % (nt) | J-GENE and allele | D-GENE and allele | AA Junction | Junction frame |
|---|---|---|---|---|---|---|---|
| $V_H$ | Musmus IGHV5-9-1*01 F | productive | 95.83% (276/288 nt) | Musmus IGHJ3*01 F | Musmus IGHD1-2*01 F | CTRHA YEAWF AYW | in-frame |
| $V_L$ | Musmus IGKV4-68*01 F | productive | 98.19% (271/276 nt) | Musmus IGKJ4*01 F | - | CQQWN SHPLTF | in-frame |

Total RNA was isolated from the hybridoma cells following the technical manual of RNeasy Plus Micro Kit (QIAGEN, Cat. No. 74034). Total RNA was then reverse-transcribed into cDNA using either isotype-specific anti-sense primers or universal primers following the technical manual of SMARTScribe Reverse Transcriptase (TaKaRa, Cat. No. 639536). Antibody fragments of heavy chain and light chain were amplified according to the standard operating procedure (SOP) of rapid amplification of cDNA ends (RACE) of GenScript. Amplified antibody fragments were cloned into a standard cloning vector separately. Colony PCR was performed to screen for clones with inserts of correct sizes.

#### 12.9. Animals and Ethical Approval

C57BL/6 mice, aged 8 weeks, were obtained from Envigo s.r.l. and housed in a specific pathogen-free (SPF) facility under controlled conditions (22 ± 2°C, 55 ± 10% humidity, 12-hour light/dark cycle). All animal procedures were approved by the Ministero della Salute (26/2014) conducted in accordance with institutional and national guidelines.

**12.10. Monoclonal Antibody Administration**

Mice were randomly assigned to treatment and control groups (n = 8 per group). The treatment group received tmCLIC1omab a dose of 10 mg/kg, administered via intravenous (i.v.) injection twice per week for a total of 3 weeks. Control animals received equivalent volumes of vehicle (isotype control antibody) on the same schedule.

**12.11. Body Weight Monitoring**

Body weight was measured twice weekly, prior to each antibody injection, using a calibrated digital scale. Baseline weight was recorded on Day 0 before the first treatment. Body weights were monitored throughout the dosing period and expressed as absolute weight (grams). Animals were observed daily for signs of stress, discomfort, or adverse effects. Any animal exhibiting a weight loss of >20% from baseline, severe behavioral changes, or other signs of distress was removed from the study and humanely euthanized according to ethical guidelines (**Fig. 6p**).
